# Effects of Developmental Lead and Phthalate Exposures on DNA Methylation in Adult Mouse Blood, Brain, and Liver Identifies Tissue- and Sex-Specific Changes with Implications for Genomic Imprinting

**DOI:** 10.1101/2023.09.29.560131

**Authors:** Rachel K. Morgan, Kai Wang, Laurie K. Svoboda, Christine A. Rygiel, Claudia Lalancette, Raymond Cavalcante, Marisa S. Bartolomei, Rexxi Prasasya, Kari Neier, Bambarendage P.U. Perera, Tamara R Jones, Justin A. Colacino, Maureen A. Sartor, Dana C. Dolinoy

## Abstract

**Background:** Maternal exposure to environmental chemicals can cause adverse health effects in offspring. Mounting evidence supports that these effects are influenced, at least in part, by epigenetic modifications.

**Objective:** We examined tissue- and sex-specific changes in DNA methylation (DNAm) associated with human-relevant lead (Pb) and di(2-ethylhexyl) phthalate (DEHP) exposure during perinatal development in cerebral cortex, blood, and liver.

**Methods:** Female mice were exposed to human relevant doses of either Pb (32ppm) via drinking water or DEHP (5 mg/kg-day) via chow for two weeks prior to mating through offspring weaning. Whole genome bisulfite sequencing (WGBS) was utilized to examine DNAm changes in offspring cortex, blood, and liver at 5 months of age. Metilene and methylSig were used to identify differentially methylated regions (DMRs). Annotatr and Chipenrich were used for genomic annotations and geneset enrichment tests of DMRs, respectively.

**Results:** The cortex contained the majority of DMRs associated with Pb (69%) and DEHP (58%) exposure. The cortex also contained the greatest degree of overlap in DMR signatures between sexes (n = 17 and 14 DMRs with Pb and DEHP exposure, respectively) and exposure types (n = 79 and 47 DMRs in males and females, respectively). In all tissues, detected DMRs were preferentially found at genomic regions associated with gene expression regulation (e.g., CpG islands and shores, 5’ UTRs, promoters, and exons). An analysis of GO terms associated with DMR-containing genes identified imprinted genes to be impacted by both Pb and DEHP exposure. Of these, *Gnas* and *Grb10* contained DMRs across tissues, sexes, and exposures. DMRs were enriched in the imprinting control regions (ICRs) of *Gnas* and *Grb10*, with 15 and 17 ICR-located DMRs across cortex, blood, and liver in each gene, respectively. The ICRs were also the location of DMRs replicated across target and surrogate tissues, suggesting epigenetic changes these regions may be potentially viable biomarkers.

**Conclusions:** We observed Pb- and DEHP-specific DNAm changes in cortex, blood, and liver, and the greatest degree of overlap in DMR signatures was seen between exposures followed by sex and tissue type. DNAm at imprinted control regions was altered by both Pb and DEHP, highlighting the susceptibility of genomic imprinting to these exposures during the perinatal window of development.

## Introduction

The health impacts of toxicant exposures during early life, such as lead (Pb) and phthalates (e.g., di(2-ethylhexyl) phthalate, DEHP) can be framed within the Developmental Origins of Health and Disease (DOHaD) hypothesis.^1^ This hypothesis postulates that exposures during sensitive periods of development alter an organism’s normal developmental programming, triggering a myriad of effects on growth and maturation that can persist into adulthood. Developmental exposures can impact gene expression long-term by altering the epigenome, which can have significant repercussions for health and disease.^2^ Epigenetics refers to mitotically heritable and potentially reversible mechanisms modulating gene expression that are independent of the DNA sequence,^3^ with the most abundantly studied mechanism being DNA methylation (DNAm). DNAm entails the addition of a methyl group to the fifth position of cytosine base adjacent to a guanine (CpG, in the majority of cases), generating what are commonly referred to as methylated cytosines (5mC), by DNA methyltransferases (DNMTs).^4^ Increased levels of 5mC within promoters and enhancers are typically associated with decreased transcription factor binding and subsequent decreases in gene expression.^5^ Patterns of 5mC undergo waves of reprogramming (i.e., global demethylation and remethylation) during critical windows of *in utero* development, making these periods susceptible targets of developmental exposures.^6^

Tight epigenetic regulation of imprinted genes is critical for early growth and development.^7,8^ Imprinted genes are expressed in a mono-allelic fashion, determined in a parent-of-origin manner. For instance, a paternally expressed gene will contain an active paternal allele and an inactive (e.g., methylated and thus imprinted) maternal allele. The DNAm patterns of imprinted genes expressed at specific developmental stages are important during growth and early development.^9,10^ Once DNAm patterns have been established for these genes, often within imprinting control regions (ICRs) in gametes, they are maintained through fertilization and extensive epigenetic reprogramming events.^11,12^ The specificity required to maintain patterns of genomic imprinting and re-establish DNAm in a parent-of-origin manner following waves of global demethylation make gestational periods particularly sensitive to environmental exposures. Environmentally-induced disruption of epigenetic processes during early development have been associated with changes in imprinted gene regulation and adverse health outcomes.^13,14^

A variety of environmental exposures, including Pb and DEHP, have been associated with altered patterns of DNAm in humans and mice.^15,16^ Pb is a known neurotoxicant, with developmental exposures linked to neurological damage and cognition deficits in early life, as well as with increased risk of degenerative neurological disease later in life.^15^ Although blood lead levels (BLLs) within the U.S. population have fallen dramatically, nearly 94% between 1976-1980 and 2015-2016, there is still concern regarding chronic low-levels of Pb exposure.^17^ This is especially true for early life exposures, as the developing brain and other organ systems are particularly susceptible to the toxic effects of Pb.^18^ Common sources of Pb exposure continue to be contaminated drinking water from leaded pipes as well as dust and chipping paint in older homes.^19,20^ Exposure to DEHP, a phthalate commonly used as a plasticizer, has become ubiquitous, with most U.S. adults having detectable levels of DEHP metabolites in their urine.^21^ DEHP is a known endocrine disruptor, with developmental exposures associated with altered metabolic function.^16,22^ Common routes of DEHP exposure include personal care products, food and beverage containers, and medical equipment, making gestational and developmental exposures common.^23^ Despite great progress over the years, gaps in knowledge remain as to whether perinatal Pb or DEHP exposure-mediated changes in DNAm have implications for long-term disease risk, whether there are sex-specific effects, and if these changes are conserved among tissues.

As a part of the Toxicant Exposures and Responses by Genomic and Epigenomic Regulators of Transcription (TaRGET II) Consortium,^24^ we utilized a mouse model of human-relevant perinatal Pb and DEHP exposures to investigate genome-wide tissue- and sex-specific associations with changes in DNAm. Whole genome bisulfite sequencing (WGBS) quantified DNAm changes in blood (an easily accessible and therefore considered a “surrogate” tissue) as well as cortex and liver (two tissues often difficult to access, representing “target” tissues) collected from male and female 5-month-old mice, with and without perinatal Pb or DEHP exposures. We assessed whether perinatal Pb- or DEHP-exposed mice displayed changes in DNAm across the genome and identified imprinted genes as a relevant gene class common to these two exposures. We additionally tested whether DNAm patterns in the surrogate tissue (blood) correlated with those seen in target tissues, to determine if blood provides a viable signature for Pb- or DEHP-induced epigenetic changes in these two tissues, and how these patterns differed between males and females.

## Methods

### Animal exposure paradigm and tissue collection

Wild-type non-agouti *a/a* mice were obtained from an over 230-generation colony of viable yellow agouti (*A^vy^*) mice, which are genetically invariant and 93% identical to the C57BL/6J strain.^25^ Virgin *a/a* females (6-8 weeks old) were randomly assigned to control, Pb-acetate water, or DEHP-chow two weeks prior to mating with virgin *a/a* males (7-9 weeks old). Pb- and DEHP-exposure were conducted *ad libitum* via distilled drinking water mixed with Pb-acetate or 7% corn oil chow mixed with DEHP. The Pb-acetate concentration was set as 32ppm to model human relevant perinatal exposure, where we have previously measured murine maternal BLLs around 16-60 ug/dL (mean: 32.1 ug/dL).^26^ DEHP was dissolved in corn oil from Envigo to create a customized stock solution, to produce 7% corn oil chow for experimentation. The DEHP exposure level was selected based on a target maternal dose of 5 mg/kg-day and assumes that a pregnant and nursing female mice weighs approximately 25 g and ingests roughly 5 g of chow per day. This target dose was selected as previous literature demonstrates obesity-related phenotypes in offspring exposed to 5 mg/kg-day DEHP during early development,^22,27^ and this dosage falls within the range of exposures previously documented in humans.^28^ All animals were maintained on a phytoestrogen-free modified AIN-93 G diet (Td.95092, 7% corn oil diet, Envigo) while housed in polycarbonate-free cages. Animal exposure to Pb or DEHP continued through gestation and lactation until weaning at post-natal day 21 (PND21) when pups were switched to either Pb-free drinking water or DEHP-free chow. Perinatal exposure, thus, occurred in offspring throughout fetal development and the first three weeks after birth. Offspring were maintained until 5 months of age. This study included n ≥ 5 males and n ≥ 5 females for Pb-exposed, DEHP-exposed, and control groups, each containing 1 male and 1 female mouse per litter; and a final samples size of n = 108 once tissues (i.e., cortex, blood, and liver) were collected. All animals and collected tissues were included in subsequent analyses, with no exclusions necessary. Prior to euthanasia, mice were fasted for 4 hours during the light cycle beginning in the morning, with euthanasia and tissue collection occurring in the afternoon. Immediately following mouse euthanasia with CO_2_ asphyxiation, blood was collected through cardiac puncture, followed by dissection of the cortex and liver, which were immediately flash frozen in liquid nitrogen and stored at −80°C. Animal collection was standardized to between 1pm to 3pm and collection order was randomized daily. For each mouse, one investigator (KN) administered the treatment and was therefore aware of the treatment group allocation. All investigators completing subsequent molecular assays were blinded to treatment group, until treatment group was analyzed during bioinformatic analyses. All mouse procedures were approved by the University of Michigan Institutional Animal Care and Use Committee (IACUC), and animals were treated humanely and with respect. All experiments were conducted according to experimental procedures outlined by the NIEHS TaRGET II Consortium.^24^ In drafting this manuscript, ARRIVE reporting guidelines were used to ensure quality and transparency of reported work.^29^

### DNA extraction and whole genome bisulfite sequencing

DNA extraction was performed using the AllPrep DNA/RNA/miRNA Universal Kit (Qiagen, Cat. #80224). Additional details about the animal exposures, blood collection, and blood DNA extraction can be found in previously published protocols.^30^ Genomic DNA (gDNA) was used in the preparation of WGBS libraries at the University of Michigan Epigenomics Core. gDNA was quantified using the Qubit BR dsDNA kit (Fisher, Cat. #Q32850), and quality assessed using Agilent’s Genomic DNA Tapestation Kit (Agilent, Cat. #A63880). For each sample, 200 ng of gDNA was spiked with 0.5% of unmethylated lambda DNA and sheared using a Covaris S220 (10% Duty Factor, 140W Peak Incident Power, 200 Cycle/Burst, 55s). A 2 µl aliquot of processed gDNA was taken to assess shearing using an Agilent High Sensitivity D1000 Kit (Agilent, Cat. #G2991AA). Once shearing was assessed, the remaining gDNA was concentrated using a Qiagen PCR Purification column and processed for end-repair and A-tailing. Ligation of cytosine-methylated adapters was done overnight at 16°C. Following this, ligation products were cleaned using AMPure XP Beads (Fisher, Cat. #NC9933872) before processing for bisulfite conversion using the Zymo EZ DNA Methylation Kit (Zymo, Cat. #D5001), and by amplifying the bisulfite converted products over 55 cycles of 95°C for 30 seconds followed by 55°C for 15 minutes, according to the manufacturer’s guidelines. After cleanup of the bisulfite converted products, final libraries were amplified over 10 cycles by PCR using KAPA Uracil+ Ready Mix (Fisher, Cat. #501965287) and NEB dual indexing primers. Final libraries were cleaned with AMPure XP beads, concentration assessed using the Qubit BR dsDNA Kit and library size assessed on the Agilent High Sensitivity D1000 Tapestation Kit. Prior to pooling, each library was quantified using KAPA Library Quantification Kit (Fisher, Cat. #501965234). We constructed four different pools of 18 libraries and each pool was sequenced on an Illumina NovaSeq6000 S4 200 cycle flow cell (PE-100) at the University of Michigan Advanced Genomics Core. Unless otherwise stated, all enzymes used in library generation were purchased from New England Biolabs. Adapters with universally methylated cytosines were synthesized by Integrated DNA Technologies (IDT).

### Data processing, quality control, and differential DNA methylation analysis

FastQC^31^ (v0.11.5) and MultiQC^32^ (v1.8) were used to assess the quality of all sequenced samples. Sequencing adapters and low-quality bases were removed by Trim Galore^33^ (v0.4.5). After trimming, reads shorter than 20 bp were removed from further analysis. Bismark^34^ (v0.19.0) with Bowtie 2^35^ (v2.3.4) as backend alignment software were used for read alignment and methylation calling with Genome Reference Consortium Mouse Build 38 (mm10) as the reference genome. All alignments were performed with 0 mismatches and multi-seed length of 20 bp. The bisulfite conversion rates were calculated through the unmethylated lambda phage DNA spike-ins. Metilene^36^ (v0.2.8) and R Bioconductor package methylSig^37^ (v1.4.0) were used to identify the differentially methylated regions (DMRs) independently. CpG sites with less than 10 reads or more than 500 reads were excluded from DMR detection. For methylSig, CpG sites that had reads covered in fewer than 4 samples within a treatment group were filtered out for DMR identification. Tiling windows were used with methylSig to identify DMRs, with a window size of 100 bp. For metilene, DMRs were identified *de novo* with at least 5 CpGs in a single DMR. For both methods, an FDR cutoff of < 0.15 and a DNAm difference of >5% were applied to select significant DMRs. All overlapping DMRs from methylSig and metilene were confirmed to be in the same direction and merged for downstream analysis (**Supplementary Table 1**). A minimum overlap cutoff of ≥ 10bp was applied to identify overlapping DMRs between tissues, sexes, and exposures, based on DMR coordinates, with no specification of methylation change direction considered for the purposes of initial comparisons. The annotatr Bioconductor package^38^ was used to annotate all significant DMRs associated with genes and genomic locations, including CpG islands, CpG shores, CpG shelves, promoters, exons, introns, 5’ UTRs, 3’ UTRs, enhancers, and regions 1-5kb upstream of transcription start sites (TSSs). Random genomic regions were generated and annotated with annotatr for each tissue using the mm10 reference genome. These random regions were used as background information to show the distribution of the genomic annotation of the DMRs if distributed purely by chance. An overview of the complete methods is illustrated in **Figure 1**.

**Figure 1:**
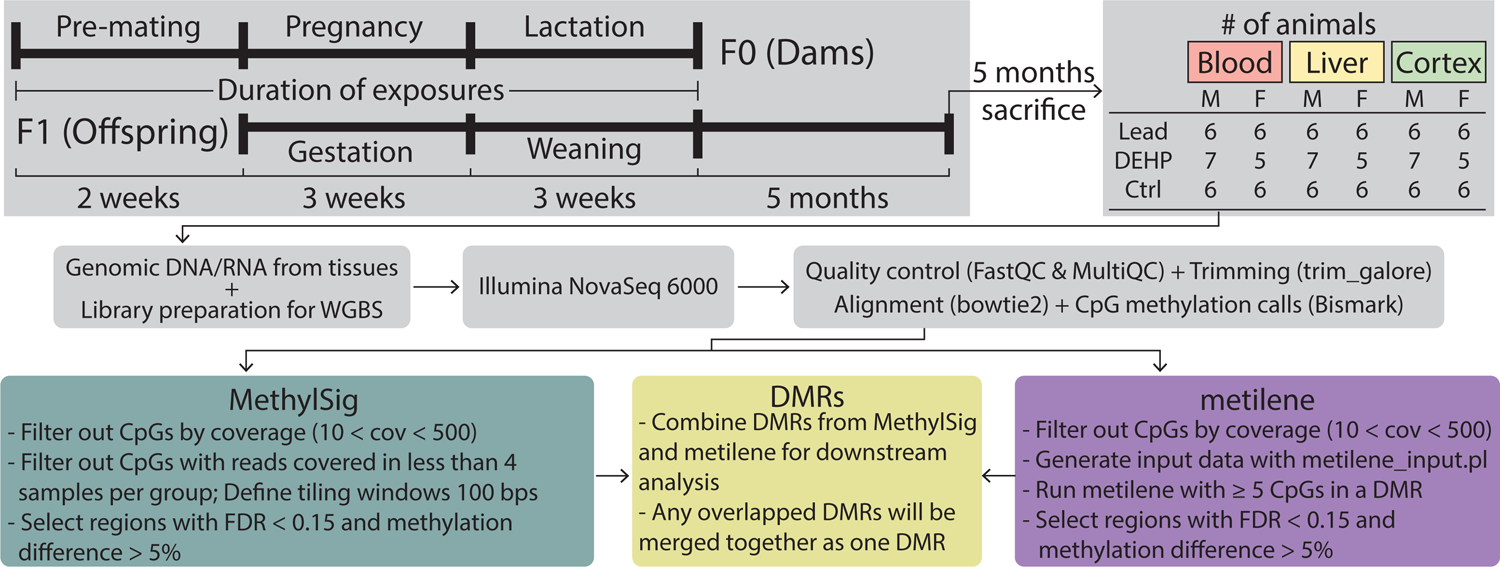
Overview of experimental workflow. F0 generation females (6-8 weeks of age) were exposed to either 32ppm of Pb via drinking water or 5mg/kg-day of DEHP via food, beginning two weeks prior to mating using virgin males (8-10 weeks of age). Exposure to Pb or DEHP or control continued through gestation and weaning, when F1 mice were removed from the dams and placed on control water or chow. At 5 months of age, F1 mice were sacrificed, and genomic DNA was extracted from blood, liver, and cortex tissues. DNA was used to prepare libraries for Whole Genome Bisulfite Sequencing (WGBS). Following initial data processing, Differentially Methylated Regions (DMRs) were called using MethylSig and metilene.

### Geneset enrichment test

R Bioconductor package Chipenrich^39^ (v2.16.0) was used to perform gene set enrichment testing of Gene Ontology (GO) terms enriched with significant DMRs. Twelve analyses were performed stratified by each tissue and sex (i.e., male cortex, male blood, male liver, female cortex, female blood, and female liver) across each exposure group (i.e., Pb, DEHP, and control). Gene assignments were determined with the *nearest_tss* locus definition in the *chipenrich* function to find all three categories of ontology (i.e., Biological Process (BP), Cellular Component (CC), and Molecular Function (MF)). An FDR cutoff of < 0.05 was applied for selecting significantly enriched GO terms. GO terms containing fewer than 15 genes or more than 500 genes were removed from analysis.

### Mouse imprinted genes and imprinted control regions

DMRs were compared to mouse imprinted genes and ICRs. We compiled a reference list of imprinted genes using previously documented efforts^40–42^ and obtained ICR coordinates from Wang et al.^43^ The valr R package^44^ (0.6.4) was used to identify overlapping regions between the DMRs and ICRs. A Binomial test was used to assess whether the DMRs were significantly enriched in ICRs and an adjusted p-value < 0.05 cutoff was utilized for identifying significant results.

## Results

### Differentially methylated regions among perinatally Pb- and DEHP-exposed tissues

Among Pb-exposed tissues, the majority of the DMRs were detected in the cortex (male (M) = 688, female (F) = 746), followed by blood (M = 243, F = 292), and liver (M = 100, F = 36). A similar pattern was observed in DEHP-exposed tissues, with the majority of DMRs detected in the cortex (M = 587, F = 661), followed by blood (M = 312, F = 477), and liver (M = 90, F = 40) (**Figure 2A**). There was limited overlap in DMRs between each tissue type, relative to the total number detected in each tissue and sex (**Figure 2B**). For instance, Pb-exposed animals had only few DMRs appear in multiple tissues. Males had 3 common DMRs among all three tissues, with 5 DMRs each overlapping between cortex and blood, between cortex and liver, and between liver and blood. Females had 7 common DMRs between cortex and blood, 3 between cortex and liver, and 1 between liver and blood, with no DMRs detected in all three tissues. Similar patterns were presented in DEHP-exposed animals, wherein males had 1 DMR common to all three tissues and 10 detected in cortex and blood, and no overlap among the remaining tissue pairs. DEHP-exposed females had more overlapping DMRs compared to males, with 2 DMR common to all tissues, 13 in both cortex and blood, 5 in cortex and liver, and 3 in liver and blood (**Figure 2B**).

**Figure 2:**
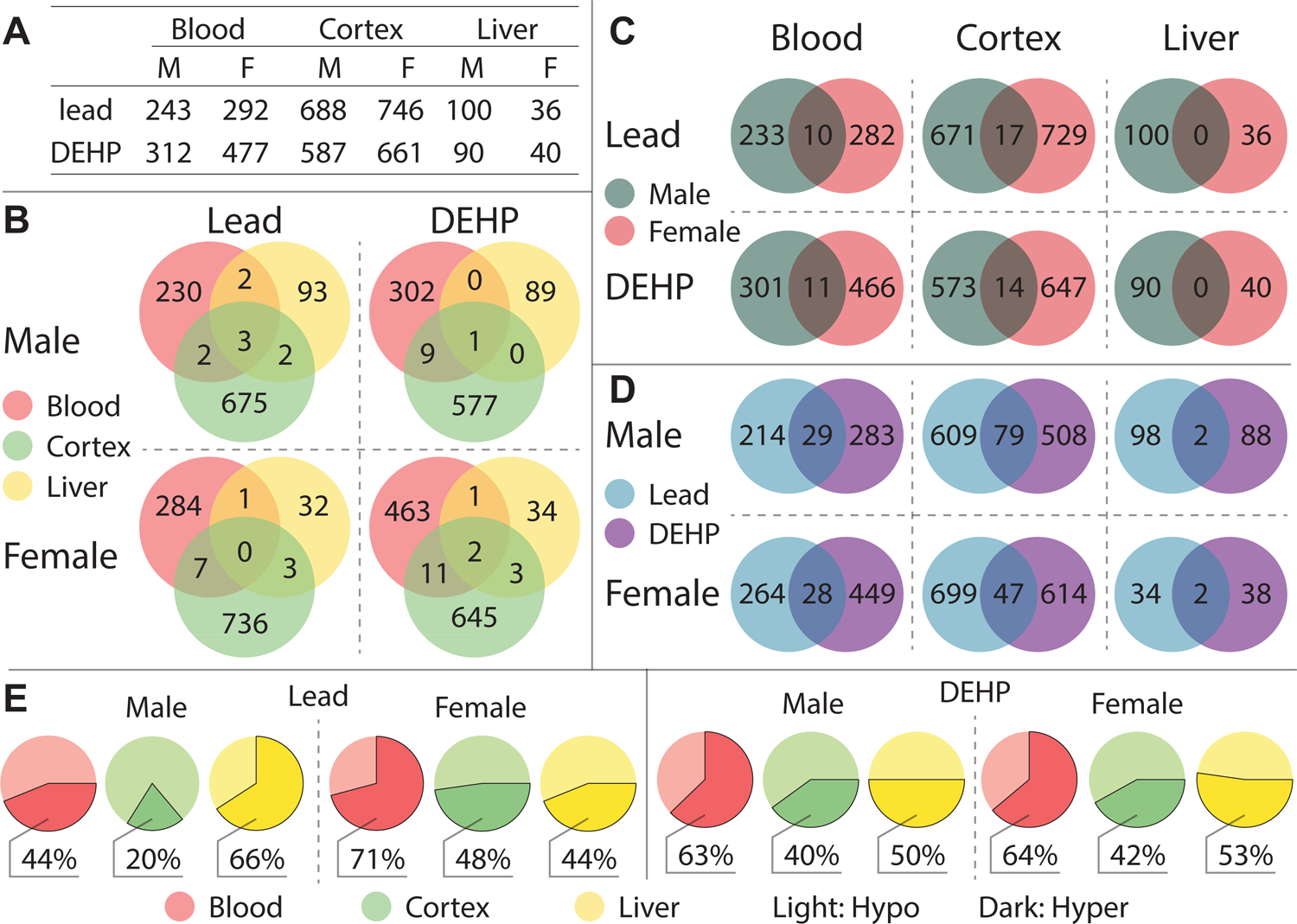
Summary of detected Differentially Methylated Regions. Differentially Methylated Regions (DMRs) were categorized by tissue (blood, cortex, and liver), sex (F: female, M: male), and exposure group (Pb, DEHP, and control) (**2A**), and DMRs found in more than one tissue type were further categorized by sex and exposure (**2B**). DMRs shared by both sexes (**2C**) and by exposure group (**2D**) were quantified and broken down by tissue type. Proportions of DMR directional changes were generally summarized for each tissue-sex-exposure combination, designated by DNA hyper (more methylated) or hypo (less methylated), in comparison to controls (**2E**).

Relative to the low overlap in exposure associated DMRs between tissues, there was more DMR similarity between the sexes when stratified by tissue (**Figure 2C**), with the exception of the liver. In Pb-exposed animals, 17 and 10 DMRs were common to both males and females in the cortex and blood, respectively. Similarly, in DEHP-exposed animals, 14 and 11 DMRs were found in both males and females in the cortex and blood, respectively (**Figure 2C**). Overall, the greatest degree of DMR overlap was found between exposure types. Pb- and DEHP-exposed cortex has the greatest degree of overlap, with 79 and 47 DMRs detected under both exposure conditions in males and females, respectively (**Figure 2D**). 29 and 28 DMRs appeared in both exposure conditions in male and female blood, respectively, whereas Pb- and DEHP-exposed liver shared 2 DMRs in each sex (**Figure 2D**).

Patterns in the direction of DNA methylation changes (DNA hyper or hypomethylation) were tissue, sex, and exposure specific (**Figure 2E**). Among Pb- and DEHP-exposed cortex, the majority of DMRs detected in males and females were hypomethylated, with slightly greater rates of hypomethylation seen in males (Pb male = 80%, Pb female = 52%, DEHP male = 60%, DEHP female = 58%). DMRs in Pb-exposed female blood, as well as DEHP-exposed male and female blood, tended to be hypermethylated (Pb female = 71%, DEHP male = 63%, DEHP female = 64%). In contrast, among Pb-exposed male blood, 56% of DMRs were hypomethylated. Patterns of directionality were more distinct between exposure types in the liver. Pb-exposed male liver presented a high proportion of hypermethylated DMRs (66%), whereas Pb-exposed female liver has slightly more hypomethylated DMRs (56%). DMR direction was roughly evenly split in DEHP-exposed liver, with 50% and 53% of DMRs hypermethylated in males and females, respectively (**Figure 2E**). **Supplementary Table 1** provides a summary of all DMRs detected in this analysis.

### Prevalence of detected DMRs in mouse genomic regions

The DMRs detected in this study occurred in specific genomic regions to a greater degree than would have been expected by a random distribution generated for comparison, given known patterns of CpG sites in the mouse genome (mm10). According to **Figure 3** and **Supplementary Table 2**, detected DMRs mapped to CpG islands to a greater degree than would have been expected by chance (3.37-19.07% of all DMRs across sex, tissues, and exposures, compared to 0.12-0.29% at random). In blood and cortex across both sexes and exposures, more DMRs were detected in 5’ UTRs than predicted 4.03-8.79%, compared to 0.18-0.4% under a random distribution), and a similar pattern was observed in liver of Pb-exposed males and females (2.02-2.91%) as well as DEHP-exposed females (4.8%, compared to 0.29% under a random distribution). Several transcriptional regulatory regions demonstrated significant derivation from what would be expected by chance as well. DMRs were present in promoter regions 2.83-6.61 times more than would have been predicted by chance across all conditions (7.3-14.41%, compared to 1.74-2.58% at random). Exons were another notable location of DMRs, with 1.76-4.99 times more DMRs than what would have been seen under a random distribution (6.74-18.65%, compared to 3.37-3.84% at random). Conversely, there were fewer DMRs detected in the open sea (11.02-21.13% in blood, 18.40-44.94% in liver, and 19.91-25.87% in cortex) than would be expected by chance (54.56-58.09%) (**Figure 3, Supplementary Table 2**).

**Figure 3:**
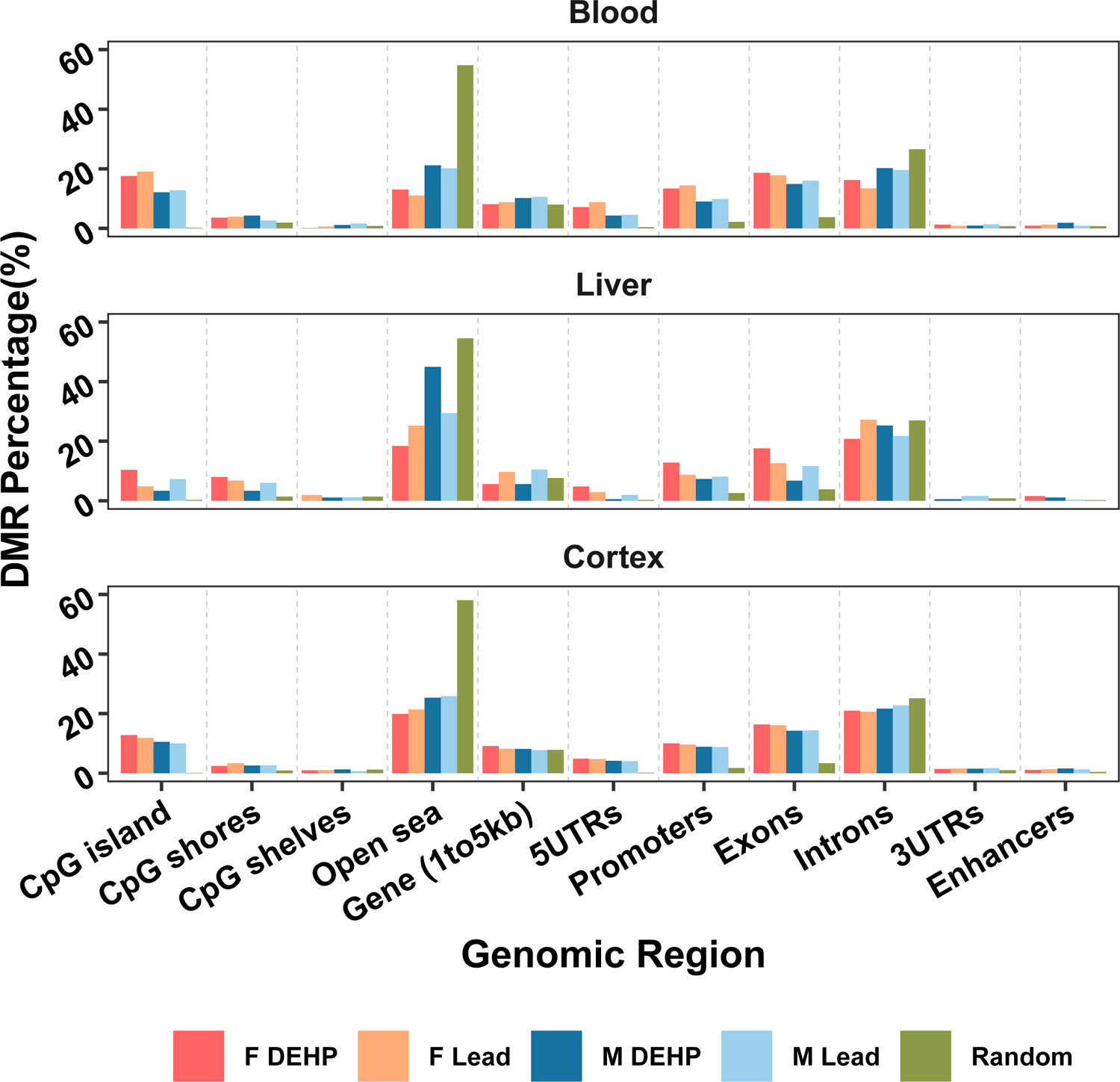
Genomic region of detected Differentially Methylated Regions. Differentially Methylated Regions (DMRs) were mapped to the mouse reference genome (mm10) and their genomic region annotated as percentage of total DMRs (comparing control and exposed samples) for that sex and exposure within each tissue. This distribution was compared to what would be expected in a random distribution.

### Gene Ontology terms associated with differentially methylated region-containing genes

DMRs were annotated using annotatr R Bioconductor package, and a summary of the overlap in DMR-containing genes across sexes, tissues, and exposures can be found in **Supplementary Figure 1**. Chipenrich was used to perform geneset enrichment tests and Gene Ontology (GO) Resource was used to identify DMR-related GO terms. The number of DMR-containing genes associated with each GO result from both Pb- and DEHP-exposed samples are summarized in **Supplementary Table 3**. Within Pb-exposed tissues, cortex had the greatest number of Gene Ontology Biological Pathway (GOBP)-related DMR-containing genes in both males (85) and females (94). DMR-associated GOBPs in female cortex were dominated by metabolic processes (35 out of 94 genes), whereas male cortex contained an abundance of DMR-containing genes related to gene expression regulation (e.g., DNA methylation or demethylation and miRNA gene silencing) (16 out of 85). The most common biological process associated with Pb exposure was genomic imprinting (GO:0071514), which appeared in male cortex, blood, and liver, as well as female cortex. In total, DMRs were detected in 21 genes associated with genomic imprinting in these tissues (**Figure 4**).

**Figure 4:**
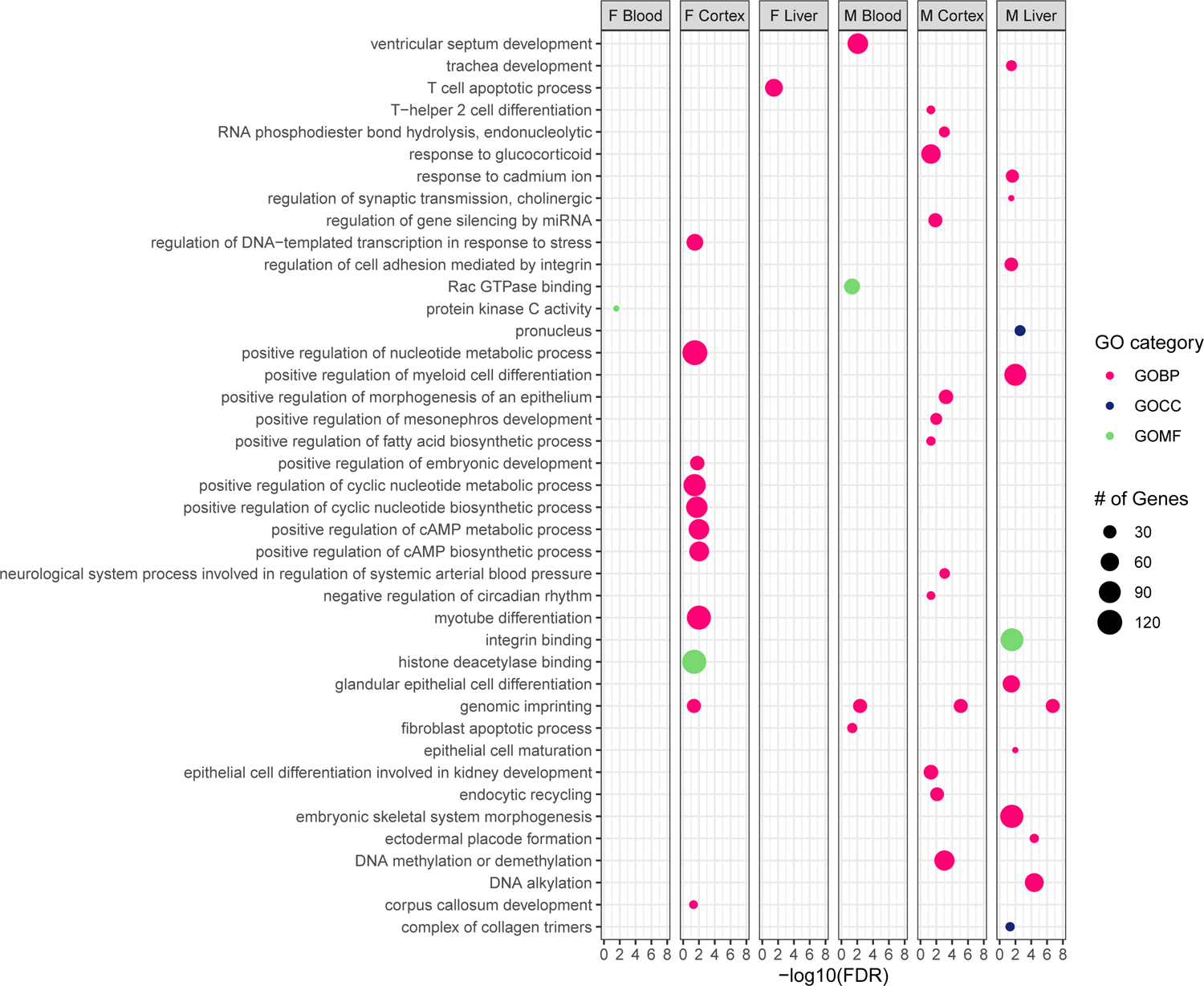
GO-terms associated with Differentially Methylated Region-containing genes among Pb-exposed tissues. Differentially Methylated Region-containing genes found in Pb-exposed tissues were submitted for Gene Ontology (GO) term analysis across three categories: Biological Process (GOBP), Cellular Component (GOCC), and Molecular Function (GOMF).

In DEHP-exposed samples, a greater number of DMR-containing genes were associated with various GO terms compared to Pb-exposed, especially the female cortex, which contained 179 genes associated with various GOBPs, most notably those associated with development (e.g., organ development, differentiation, and morphogenesis) (148 of 179). Male cortex contained far fewer GO term-associated DMRs compared to females (66 compared to 179), and there was an abundance of genes associated with gene expression regulation (10) and cellular organization (20). As with Pb-exposed tissues, the only GO term common to more than one tissue-sex combination among DEHP samples was genomic imprinting, which was associated with DMRs in 9 genes across male blood and cortex (**Figure 5**).

**Figure 5:**
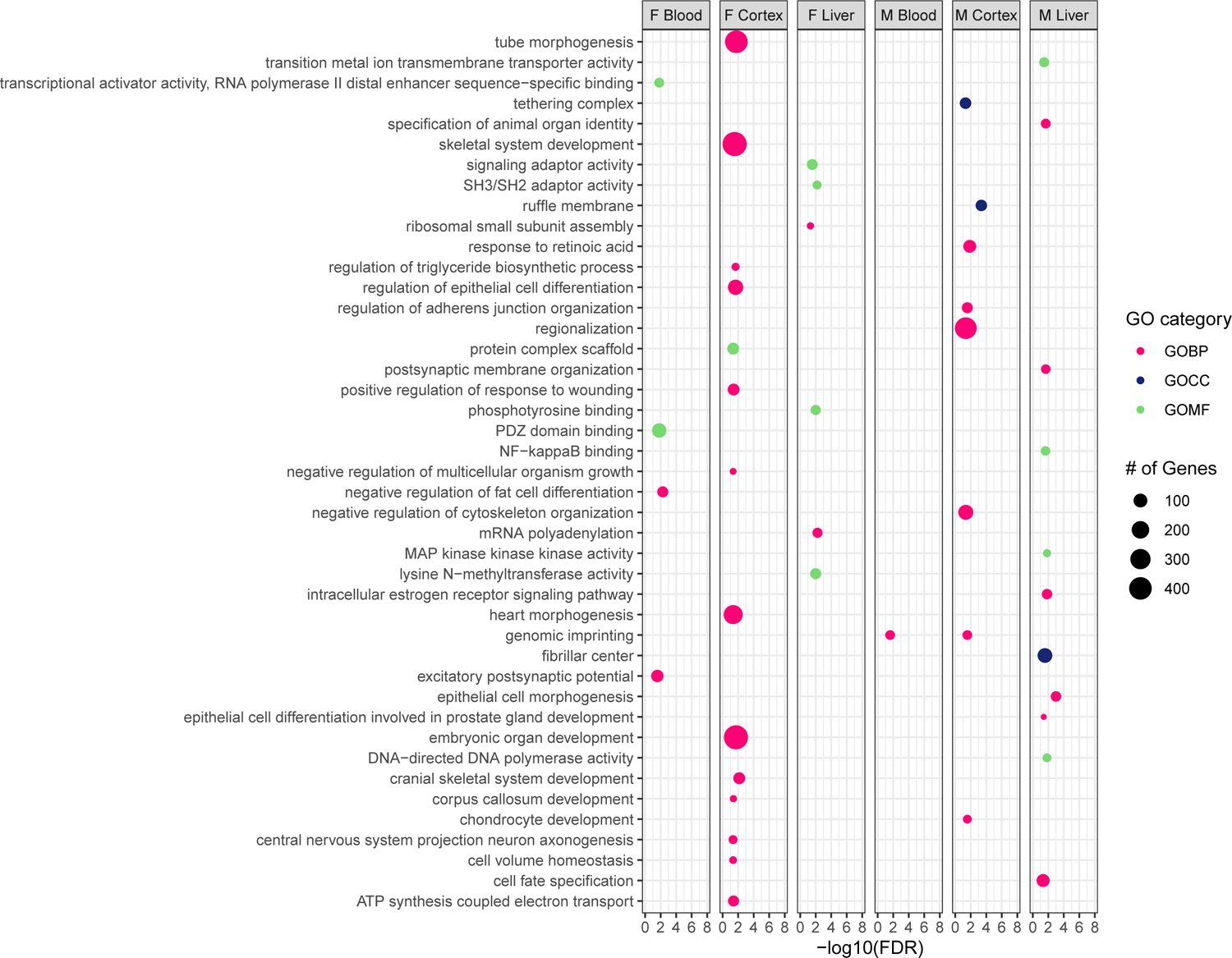
GO-terms associated with Differentially Methylated Region-containing genes among DEHP-exposed tissues. Differentially Methylated Region-containing genes found in DEHP-exposed tissues were submitted for Gene Ontology (GO) term analysis across three categories: Biological Process (GOBP), Cellular Component (GOCC), and Molecular Function (GOMF).

### DNA methylation changes at imprinted loci

The appearance of imprinted genes in both exposure models during pathway analysis (**Figures 4 and 5**) was motivation to take a closer look at the effects of Pb and DEHP exposure on imprinted genes. All tissue types, across both sexes and exposures had detectable changes in DNAm within imprinted genes (**Supplementary Figures 2-5**). A reference list of imprinted genes used in this analysis can be found in **Supplemental Table 4**, and genes that did not contain a DMR in any tissue were omitted from the final figure. Cortex had the greatest number of DMRs as well as the greatest magnitude of methylation changes in assessed imprinted genes. 73 Pb-associated DMRs were detected in cortex at imprinted genes (46 in males and 27 in females with magnitude changes of 5.03-23.77%) and 67 were detected in DEHP-exposed cortex (37 in males and 30 in females with magnitude changes of 5.2-24.9%). 36 Pb-associated DMRs were detected in blood at imprinted genes (16 in males and 20 in females with magnitude changes of 5.04-20.1%) and 55 were detected in DEHP-exposed blood (32 in males and 23 in females with magnitude changes of 5.4-28.4%). Liver contained fewer changes in DNAm at imprinted genes, compared to blood and cortex, for each sex-exposure combination, with 10 DMRs in Pb-exposed liver (9 in males and 1 in females with magnitude changes of 6.8-19.4%) and 11 DMRs in DEHP-exposed liver (3 in males and 8 in females with magnitude changes of 8.8-16.3%). Blood from Pb-exposed females largely contained hypermethylated sites at imprinted genes (15/20 DMRs), while cortex from the same animals was largely hypomethylated in the same gene class (20/27 DMRs). A similar pattern was seen in DEHP-male tissues, with the bulk of detectable changes found in the blood and cortex, with the former being largely hypermethylated (29/32 DMRs in blood) and the latter hypomethylated 23/37 DMRs in cortex) (**Supplementary Table 5**).

Two imprinted genes, *Gnas* and *Grb10*, contained a notable number of exposure associated DMRs. A complete overview of these DMRs is summarized in **Figure 6** and **Supplementary Table 6**. Among Pb-exposed samples, 60% and 75% of DMRs in the *Gnas* locus were hypomethylated in males and females, respectively. In Pb-exposed blood, DMRs within the *Gnas* locus were entirely hypermethylated in females (1/1) and hypomethylated in males (3/3). In Pb-exposed liver, *Gnas* DMRs were hypermethylated (2/2). Among Pb-exposed cortex, DMRs within the *Grb10* locus were largely hypermethylated in males (66%) and hypomethylated in females (66%). A similar pattern presented in Pb-exposed blood, wherein the entirety of *Grb10* DMRs in males were hypermethylated (2/2), whereas those in females were hypomethylated (1/1). Male liver contained only hypermethylated sites (2/2) within the *Grb10* locus.

**Figure 6:**
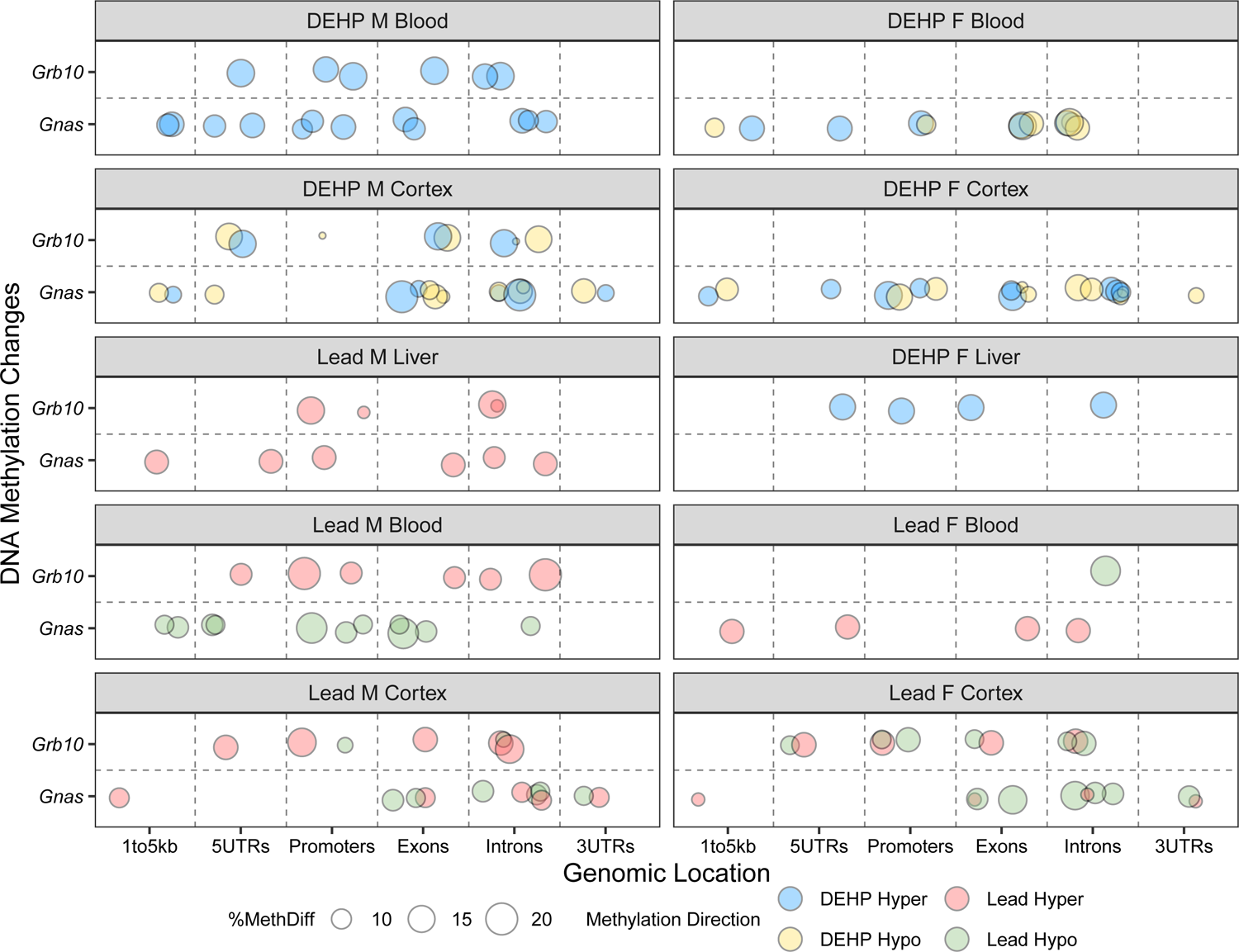
Genomic location and direction of Pb and DEHP-associated Differentially Methylated Regions in the *Gnas* and *Grb10* loci. Differentially Methylated Regions (DMRs) detected in the *Gnas* and *Grb10* loci were classified as to their genomic location within each gene. Percent change in methylation is denoted by size and direction of methylation change by color (blue = hypermethylated DMRs among DEHP samples, yellow = hypomethylation among DEHP samples, red = hypermethylation among Pb samples, green = among hypomethylation among Pb samples).

DEHP exposure was associated with more hypomethylation at the *Gnas* locus in male cortex (80%) than in females (50%). In blood, DEHP exposure associated with more hypomethylation in females (75%) but DMRs associated with this exposure in male blood were entirely hypermethylated (3/3). Regarding *Grb10*, 2/3 DMRs identified in male cortex were hypomethylated whereas 2/2 identified in male blood were hypermethylated. One hypermethylated DMR was detected in *Grb10* in DEHP-exposed female liver.

### Exposure-associated changes in imprinting control regions

Imprinted genes are regulated in part through imprinting control regions (ICRs), which are elements whose methylation is set up in the germline and that regulate gene expression and subsequent functions of imprinted gene clusters.^36^ Changes in the DNAm status of these regions can impact the expression of imprinted and non-imprinted genes within a given cluster, thus magnifying the regulatory effects of what would otherwise be a single-gene effect.^37^ *Gnas* contains two ICRs, the *Gnas* ICR and the *Nespas* ICR, while *Grb10* contains one ICR.^36,38^ The current analysis identified multiple DMRs within the ICRs of both *Gnas* (7 in ICR *Gnas* and 8 in ICR *Nespas*) and *Grb10* (17 in the *Grb10* ICR) across exposure and tissue types (**Figure 7**). A binomial test was conducted to assess whether exposure-associated DMRs occurred in these ICRs to a greater degree than would have been expected by random change. Both the *Gnas* and *Grb10* ICRs contained more DMRs than would have been expected by chance in multiple sex-exposure-tissue combinations. A summary of these findings can be found in **Supplementary Table 6**.

**Figure 7:**
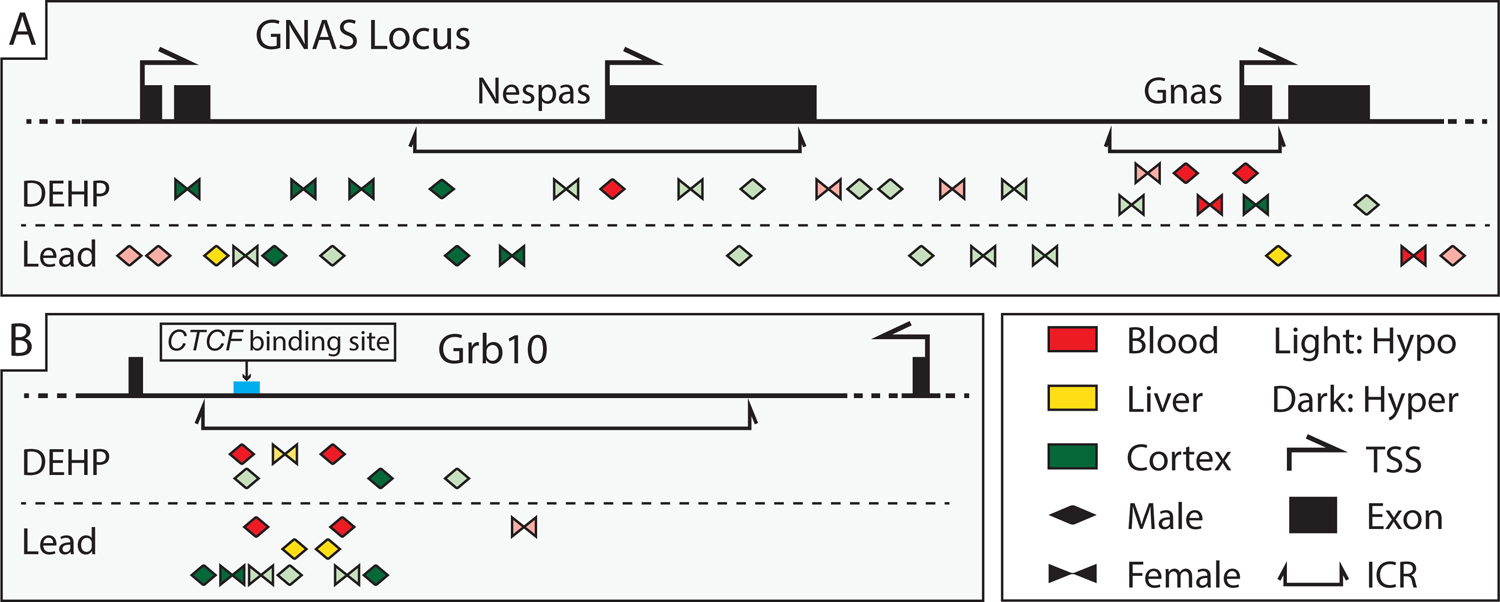
Differentially Methylated Regions detected within *Gnas* and *Grb10* Imprinting Control Regions (ICRs) among Pb and DEHP exposed tissues. (A) Differentially Methylated Regions (DMRs) overlap with *Gnas*. (B) DMRs overlap with *Grb10*. DMRs only represents the related genomic locations corresponding to the genomic coordinates of ICRs. The genomic coordinates of these DMRs can be found in Supplementary Table 4.

Pb exposure was associated with relatively limited changes in DNAm in *Gnas* ICRs when compared to *Grb10*. In the *Nespas* ICR, Pb exposure was associated with hypermethylation in female cortex (1/1 DMR) and a mix of hyper-(1/2 DMRs) and hypomethylation (1/2 DMRs) in male cortex. In the *Gnas* ICR, Pb exposure was associated only with hypermethylation in male liver (1/1 DMR) (**Figure 7A** and **Supplementary Table 7**). In the *Grb10* ICR, Pb exposure was associated again with an equal amount of hyper-(3/6) and hypomethylated (3/6) DMRs, in both male and female cortex. Pb exposure was entirely associated with hypermethylation in both male blood (2/2 DMRs) and liver (2/2 DMRs) but was associated with hypomethylation in female blood (1/1 DMR) (**Figure 7B** and **Supplementary Table 7**).

There were comparatively more changes in DNAm in the *Gnas* ICRs associated with DEHP exposure. In male cortex there was again a mix of hyper-(1/2) and hypomethylated (1/2) DMRs in the *Nespas* ICR. Unlike Pb exposure, DEHP was associated only with hypomethylated DMRs (2/2) in female cortex in the *Nespas* ICR. In male blood there was 1 and 2 hypermethylated DEHP-associated DMRs within the *Nespas* and *Gnas* ICRs, respectively. Female cortex and blood both contained a mix of hyper-(1/2) and hypomethylated (1/2) DMRs in the *Gnas* ICR associated with DEHP exposure (**Figure 7A** and **Supplementary Table 8**). Within the *Grb10* ICR, DEHP exposure was associated with a mix of hyper-(1/3) and hypomethylated (2/3) DMRs in male cortex, hypermethylated (2/2) DMRs in male blood, and 1 hypermethylated DMR in female liver (**Figure 7B** and **Supplementary Table 8**).

## Discussion

Toxicant exposures that occur during critical periods of development can have ramifications for health and well-being throughout the life-course.^45^ Perinatal Pb and DEHP exposures have been linked to aberrant brain development and metabolic function, respectively, at environmentally relevant doses.^46,47^ With regard to epigenetic mechanisms governing gene expression, Pb and DEHP exposures have both been associated with differential DNAm in human populations.^48,49^ Concurrently, it is unknown if toxicant-induced changes in difficult-to-access tissues, such as brain and liver, are reflected in more easily accessible (surrogate) tissues, such as blood. It is therefore pertinent to examine how two prominent developmental exposures, Pb and DEHP, affect gene regulation by DNAm in these target and surrogate tissues in order to assess whether DNAm could be used as a potential biomarker of changes in more difficult to access tissues, as is being evaluated in the TaRGET II Consortium.^24^

### Pb and DEHP Exposures are Associated with Sex, Tissue, and Exposure-Specific General Changes in DNA Methylation

Overall, Pb and DEHP exposures resulted in similar number of DMRs between the sexes for each of the three tissues assessed (**Figure 2A**). The cortex contained the greatest number of DMRs for each exposure, followed by blood and liver. Between the sexes, females had more DMRs across both exposures in cortex and blood, while males had more DMRs in the liver (**Figure 2A**). This overall DNAm pattern is consistent with previous reports, which showed significant changes in DNAm in female brain following gestational Pb exposure as well as in male liver following DEHP exposure.^50,51^ There was minimal overlap in DMRs between either cortex-blood (0.5-1.2% of total DMRs detected in these tissues) or liver-blood (0.25-1.5% of total DMRs detected in these tissues) (**Figure 2B**). The largest degree in DMR similarity between target-surrogate tissues was in DEHP-exposed female cortex and blood (13 similar DMRs, 1.16% of all DMRs in those tissues), followed by DEHP-exposed male cortex and blood (10 similar DMRs, 1.12% of all DMRs in those tissues). These findings suggest limited general overlap in DNAm changes across surrogate and target tissues when stratified by sex and exposure.

When the similarity of DMR signatures between the sexes was assessed for Pb and DEHP exposures, the greatest number of shared DMRs was seen in the cortex, followed by blood, with no common DMRs in the liver (**Figure 2C)**. The number of DMRs in common between the sexes did not exceed 2% of the total DMRs detected in any tissue-exposure combination. These findings highlight the need to evaluate sex-specific effects in toxicoepigenetic studies.^52,53^ A greater degree of DMR similarity was seen between exposure types, with 1-7% of total DMRs appearing in both Pb and DEHP-exposed tissues, depending on the sex and tissue (**Figure 2D)**. General trends in DMR directionality were not conserved across tissue types, adding complexity to comparisons of changes in DNAm patterns between target and surrogate tissues (**Figure 2E**). As expected, many DMRs were located in CpG islands, areas of dynamic DNAm-directed gene expression regulation.^54^ Gene promoters and exons also contained more DMRs than would have been predicted by chance (**Figure 3**).

### Exposure-Associated DMRs Occur to a Notable Degree in Imprinted Genes

An analysis of GO terms associated with DMR-containing genes identified genomic imprinting as a common category across most tissues in both sexes and exposure types (**Figures 4 and 5**). Imprinted genes are an important class with regard to early growth and development, and their epigenetically-controlled mono-allelic parent of origin nature of expression may confer particular susceptibility to the impacts of environmental exposures.^55,56^ Early disruption of imprinted gene expression and function can result in developmental disorders (e.g., pseudohypoparathyroidism type 1B and Silver-Russell syndrome, for which perturbations in gene expression regulation of *Gnas* and *Grb10*, respectively, have been implicated.^57,58^ Additionally, changes in the DNAm status of several imprinted genes have been associated with chronic conditions such as diabetes, cardiovascular disease, and cancer.^59–61^ The DNAm and hydroxymethylation status of imprinted genes is particularly susceptible to environmental exposures during early development, including Pb and DEHP.^62–64^ Epidemiological studies have linked early life Pb exposure to altered methylation in imprinted genes including insulin-like growth factor 2 (*IGF2*), which is involved in some cases of Beckwith-Wiedemann Syndrome and Silver-Russell Syndrome and maternally expressed gene 3 (*MEG3*), which is implicated in Temple syndrome and Kagami-Ogata syndrome.).^65,66^

### Imprinting Control Regions Contain Exposure- and Tissue-Specific Changes in DNA Methylation

ICRs are environmentally sensitive regulatory regions, and changes to their DNAm status can have consequences for a cluster of imprinted genes.^67^ The ICRs of both *Gnas* and *Grb10* contained numerous Pb- and DEHP-associated DMRs, with *Gnas* ICR DMRs appearing largely in the cortex and to be more prevalent with DEHP exposure, while the *Grb10* ICR contained about twice as many Pb-associated DMRs than DEHP and with much more even distribution across the studied tissues.

The *Grb10* ICR contained DMRs across all three tissues examined, with a specific DMR replicated in Pb-exposed male liver and blood. There was an additional DMR in common in DEHP-exposed male cortex and blood, but they differed in directionality (cortex = hypomethylated, blood = hypermethylated). The current study is one the few reports that examine the effects of environmental exposures on the *Grb10* ICR, with a previous report highlighting the effects on hydroxymethylation,^64^ though many more exist pertaining to changes throughout the gene.^68,69^ Much of the published work is restricted to germ cells, and so additional work is needed to assess whether *Grb10* regulation and function are impacted by the environment in the soma.

### Differential Methylation of Gnas and Grb10 Occurred in Gene Expression Regulatory Regions

*Gnas* encodes for the G-protein alpha-subunit protein, which contributes to signal transduction via cAMP generation^70^, and its imprinting dysregulation has been associated with increased insulin sensitivity, neural tube defects, and hypothyroidism.^71,72^ The imprinted expression of *Gnas* is complex, as this gene gives rise to several maternal- and paternal-specific gene products, and these patterns of expression are highly tissue-specific in mice and humans.^73–75^ In this work, *Gnas* contained a mix of hyper- and hypomethylated DMRs in the cortex, under both exposure conditions (**Figure 6**), making the prediction of the observed sustained DNAm effects at 5 months difficult to ascertain. However, given the importance of maintained imprinted expression of this locus and its various gene products in the brain, continued evaluation of the effects of exposure-induced changes in DNAm at this locus would help elucidate the functional impacts on gene product expression and subsequent physiological effects. Changes in DNAm within *Gnas* were much more uniform in blood and liver, where biallelic expression is considered to be the norm in adult mice.^70^ Distinct differences in *Gnas* DMR direction appeared between the sexes in this study (**Figure 6**). In Pb-exposed blood, *Gnas* DMRs were entirely hypomethylated in males and hypermethylated in females. Within DEHP-exposed blood, *Gnas* DMRs were entirely hypermethylated in males and a mix of hyper- and hypomethylated in females (**Figure 6**). As this work found hypomethylation in *Gnas* promoters of DEHP-exposed female cortex and blood, it would be pertinent to expand this work to additional tissues such as the thyroid to ascertain whether this relationship is consistent in an organ known to be significantly impacted by developmental changes in *Gnas* DNAm status.

*Grb10* encodes for an insulin receptor-binding protein involved in growth and insulin response and is imprinted in a tissue- and cell-type specific manner.^76,77^ This is especially true during development, as changes in *Grb10* expression across time are tissue-specific. For example, in the brain, *Grb10* imprinting status is cell-type specific during development until adulthood.^78^ There were several DMRs detected in *Grb10* in DEHP-exposed male cortex, as well as Pb-exposed male and female cortex. *Grb10* methylation appears to be cell-type specific during early brain development, with paternal expression in cortical neurons and maternal expression in glial cells.^77^ While this study was unable to assess cell-type specific changes in DNAm within the cortex, future single-cell analyses could help determine whether exposure-associated DMRs are specific to certain cellular populations. *Grb10* expression also changes significantly in the liver during development, as maternal expression is high during fetal development, but nearly all *Grb10* expression is silenced in the liver in adulthood.^79,80^ Many of the DMRs seen in *Grb10* in the liver were hypermethylated, suggesting these exposures may not result in the reactivation of this gene in adulthood, but, alternatively, may reinforce its suppression through supplemental methylation. Whether this trend was present during early development, when imprinted expression is the norm and whether that had any deleterious effects on liver development, remains to be seen. Pb exposure, on the other hand, was associated with hypomethylation of *Grb10* in female blood, another tissue in which *Grb10* is thought to be maternally expressed during early development and completely repressed during adulthood,^80^ meaning that exposure may be related to reactivation of this gene during an inappropriate time point. Future evaluation of the impact of *Grb10* expression in blood during adulthood would contribute to our understanding of the potential functional impact of this change in methylation. *Grb10* is initially expressed from the maternal allele in somatic lineages and exclusively in neurons, switches to paternal-specific expression from an alternate promoter.^77^

### Gnas and Grb10 Provide Evidence of DNA Methylation Signatures in Target-Surrogate Tissue Pairs

The ICRs of both *Gnas* and *Grb10* displayed some changes in DNAm that were replicated in both target and surrogate tissues, suggesting these regulatory regions may be of significance when attempting to identify DNAm-related biomarkers of exposure (**Figure 7**). Among Pb-exposed samples, the *Grb10* ICR contained hypermethylated DMRs in male cortex, liver, and blood, suggesting that, for this exposure, the *Grb10* ICR may be a potential region to consider when exploring male-specific DNAm biomarkers of exposure. Among DEHP-exposed samples, the *Gnas* ICR contained hyper- and hypomethylated DMRs that were seen in female cortex and blood, while the *Nespas* ICR was the location of hypermethylated DMRs in male cortex and blood. These findings suggest there may be ICR- and sex-specificity in terms of DNAm biomarkers of DEHP exposure, and that they may be particularly applicable to the cortex and blood. DEHP-associated hypermethylated DMRs were also replicated in male cortex and blood within the *Grb10* ICR, suggesting this regulatory region may be an additional candidate as a DNAm biomarker for DEHP exposure.

## Limitations

DNAm patterns vary across cell types within a given tissue.^81,82^ This study was unable to account for cell type and therefore, changes in DNAm as the result of Pb or DEHP exposure may be due to exposure-induced changes in cell type proportions.^83^ Additionally, we were not able to evaluate changes in DNA hydroxymethylation (5hmC) in these samples. This study was conducted using bisulfite conversion, which accounts for both 5mC and 5hmC, and the resulting data is unable to differentiate between these two signatures.^64^. Imprinted genes are typically 50% methylated (accounting for mono-allelic expression or repression), and this data represents DNAm averages for both alleles. Thus, any allele-specific changes in DNAm associated with Pb or DEHP cannot be detected.

## Conclusion

This study systematically evaluated changes in DNAm for cortex, blood, and liver collected from mice at 5 months-of-age following developmental exposure to either Pb or DEHP. Pb- and DEHP-specific DNAm changes were observed via DMRs, with the greatest DMR similarity seen between exposure types, with less overlap between the sexes and tissues. Genomic imprinting was impacted by Pb and DEHP exposure, as determined by GO term analysis, and imprinted genes *Gnas* and *Grb10* indicated changes in DNAm at their respective ICRs. These results indicate that imprinted gene methylation can be dysregulated by developmental environmental exposures such as Pb and DEHP and that ICRs may be useful candidates when exploring DNAm-based biomarkers of environmental exposures.

## Acknowledgements

We would like to acknowledge members of the University of Michigan Epigenomics Core and the Advanced Genomics Core, as well as the Michigan Lifestage Environmental Exposures and Disease Center (M-LEEaD) which facilitated the generation and analysis of WGBS data.

## Data Sharing

WGBS data will be uploaded to GEO. Additional data that support the findings of this study are available from the corresponding author, DCD, upon reasonable request.

## Approval for Animal Use

Work outlined in this manuscript was approved by the University of Michigan Institutional Animal Care and Use Committee (IACUC) and conducted in accordance with the highest animal welfare standards.

## References

1 Gillman MW. Developmental origins of health and disease. The New England Journal of Medicine 2005;353:1848.

2 Bernal AJ, Jirtle RL. Epigenomic disruption: the effects of early developmental exposures. Birth Defects Research Part A: Clinical and Molecular Teratology 2010;88:938–44.

3 Bollati V, Baccarelli A. Environmental epigenetics. Heredity 2010;105:105–12.

4 Lyko F. The DNA methyltransferase family: a versatile toolkit for epigenetic regulation. Nature Reviews Genetics 2018;19:81–92.

5 Siegfried Z, Simon I. DNA methylation and gene expression. Wiley Interdisciplinary Reviews: Systems Biology and Medicine 2010;2:362–71.

6 Zeng Y, Chen T. DNA methylation reprogramming during mammalian development. Genes 2019;10:257.

7 SanMiguel JM, Bartolomei MS. DNA methylation dynamics of genomic imprinting in mouse development. Biol Reprod 2018;99:252–62. 10.1093/biolre/ioy036.

8 Tucci V, Isles AR, Kelsey G, Ferguson-Smith AC, Bartolomei MS, Benvenisty N, et al. Genomic imprinting and physiological processes in mammals. Cell 2019;176:952–65.

9 Piedrahita JA. The role of imprinted genes in fetal growth abnormalities. Birth Defects Res A Clin Mol Teratol 2011;91:682–92. 10.1002/bdra.20795.

10 Moore GE, Ishida M, Demetriou C, Al-Olabi L, Leon LJ, Thomas AC, et al. The role and interaction of imprinted genes in human fetal growth. Philos Trans R Soc Lond B Biol Sci 2015;370:20140074. 10.1098/rstb.2014.0074.

11 Jima DD, Skaar DA, Planchart A, Motsinger-Reif A, Cevik SE, Park SS, et al. Genomic map of candidate human imprint control regions: the imprintome. Epigenetics 2022:1–24.

12. Horsthemke B. Mechanisms of Imprint Dysregulation.

13 Faulk C, Dolinoy DC. Timing is everything: the when and how of environmentally induced changes in the epigenome of animals. Epigenetics 2011;6:791–7.

14 Angers B, Castonguay E, Massicotte R. Environmentally induced phenotypes and DNA methylation: how to deal with unpredictable conditions until the next generation and after. Molecular Ecology 2010;19:1283–95.

15 Senut M-C, Cingolani P, Sen A, Kruger A, Shaik A, Hirsch H, et al. Epigenetics of early-life lead exposure and effects on brain development. Epigenomics 2012;4:665–74. 10.2217/epi.12.58.

16 Parsanathan R, Karundevi B. Phthalate exposure in utero causes epigenetic changes and impairs insulin signalling. Journal of Endocrinology 2014;223:47–66. 10.1530/JOE-14-0111.

17 Dignam T, Kaufmann RB, LeStourgeon L, Brown MJ. Control of Lead Sources in the United States, 1970-2017: Public Health Progress and Current Challenges to Eliminating Lead Exposure. J Public Health Manag Pract 2019;25:S13–22. 10.1097/PHH.0000000000000889.

18 Zhou F, Yin G, Gao Y, Liu D, Xie J, Ouyang L, et al. Toxicity assessment due to prenatal and lactational exposure to lead, cadmium and mercury mixtures. Environ Int 2019;133:105192. 10.1016/j.envint.2019.105192.

19 Kamenov GD, Swaringen BF, Cornwell DA, McTigue NE, Roberts SM, Bonzongo J-CJ. High-precision Pb isotopes of drinking water lead pipes: Implications for human exposure to industrial Pb in the United States. Sci Total Environ 2023;871:162067. 10.1016/j.scitotenv.2023.162067.

20 Dietrich M, Barlow CF, Entwistle JA, Meza-Figueroa D, Dong C, Gunkel-Grillon P, et al. Predictive modeling of indoor dust lead concentrations: Sources, risks, and benefits of intervention. Environ Pollut 2023;319:121039. 10.1016/j.envpol.2023.121039.

21 Wang Y, Qian H. Phthalates and Their Impacts on Human Health. Healthcare (Basel) 2021;9:603. 10.3390/healthcare9050603.

22 Lin Y, Wei J, Li Y, Chen J, Zhou Z, Song L, et al. Developmental exposure to di(2-ethylhexyl) phthalate impairs endocrine pancreas and leads to long-term adverse effects on glucose homeostasis in the rat. American Journal of Physiology-Endocrinology and Metabolism 2011;301:E527–38. 10.1152/ajpendo.00233.2011.

23 Erythropel HC, Maric M, Nicell JA, Leask RL, Yargeau V. Leaching of the plasticizer di(2-ethylhexyl)phthalate (DEHP) from plastic containers and the question of human exposure. Appl Microbiol Biotechnol 2014;98:9967–81. 10.1007/s00253-014-6183-8.

24 Wang T, Pehrsson EC, Purushotham D, Li D, Zhuo X, Zhang B, et al. The NIEHS TaRGET II Consortium and environmental epigenomics. Nature Biotechnology 2018;36:225–7.

25 Dou JF, Farooqui Z, Faulk CD, Barks AK, Jones T, Dolinoy DC, et al. Perinatal Lead (Pb) Exposure and Cortical Neuron-Specific DNA Methylation in Male Mice. Genes (Basel) 2019;10:E274. 10.3390/genes10040274.

26 Faulk C, Barks A, Liu K, Goodrich JM, Dolinoy DC. Early-life lead exposure results in dose- and sex-specific effects on weight and epigenetic gene regulation in weanling mice. Epigenomics 2013;5:487–500. 10.2217/epi.13.49.

27 Schmidt J-S, Schaedlich K, Fiandanese N, Pocar P, Fischer B. Effects of di(2-ethylhexyl) phthalate (DEHP) on female fertility and adipogenesis in C3H/N mice. Environ Health Perspect 2012;120:1123–9. 10.1289/ehp.1104016.

28 Neier K, Cheatham D, Bedrosian LD, Dolinoy DC. Perinatal exposures to phthalates and phthalate mixtures result in sex-specific effects on body weight, organ weights and intracisternal A-particle (IAP) DNA methylation in weanling mice. J Dev Orig Health Dis 2019;10:176–87. 10.1017/S2040174418000430.

29 Percie du Sert N, Hurst V, Ahluwalia A, Alam S, Avey MT, Baker M, et al. The ARRIVE guidelines 2.0: Updated guidelines for reporting animal research. PLoS Biol 2020;18:e3000410. 10.1371/journal.pbio.3000410.

30 Svoboda LK, Neier K, Wang K, Cavalcante RG, Rygiel CA, Tsai Z, et al. Tissue and sex-specific programming of dna methylation by perinatal lead exposure: implications for environmental epigenetics studies. Epigenetics 2021;16:1102–22. 10.1080/15592294.2020.1841872.

31 Andrews. FastQC A Quality Control tool for High Throughput Sequence Data. 2010. URL: https://www.bioinformatics.babraham.ac.uk/projects/fastqc/ (Accessed 15 February 2023).

32 Ewels P, Magnusson M, Lundin S, Käller M. MultiQC: summarize analysis results for multiple tools and samples in a single report. Bioinformatics 2016;32:3047–8. 10.1093/bioinformatics/btw354.

33 Krueger F. Trim Galore. 2015. URL: https://www.bioinformatics.babraham.ac.uk/projects/trim_galore/ (Accessed 15 February 2023).

34 Krueger F, Andrews SR. Bismark: a flexible aligner and methylation caller for Bisulfite-Seq applications. Bioinformatics 2011;27:1571–2. 10.1093/bioinformatics/btr167.

35 Langmead B, Salzberg SL. Fast gapped-read alignment with Bowtie 2. Nat Methods 2012;9:357–9. 10.1038/nmeth.1923.

36 Jühling F, Kretzmer H, Bernhart SH, Otto C, Stadler PF, Hoffmann S. metilene: fast and sensitive calling of differentially methylated regions from bisulfite sequencing data. Genome Res 2016;26:256–62. 10.1101/gr.196394.115.

37 Park Y, Figueroa ME, Rozek LS, Sartor MA. MethylSig: a whole genome DNA methylation analysis pipeline. Bioinformatics 2014;30:2414–22. 10.1093/bioinformatics/btu339.

38 Cavalcante RG, Sartor MA. annotatr: genomic regions in context. Bioinformatics 2017;33:2381–3. 10.1093/bioinformatics/btx183.

39 Welch RP, Lee C, Imbriano PM, Patil S, Weymouth TE, Smith RA, et al. ChIP-Enrich: gene set enrichment testing for ChIP-seq data. Nucleic Acids Res 2014;42:e105. 10.1093/nar/gku463.

40 Williamson CM, Blake A, Thomas S, Beechey CV, Hancock J, Cattanach BM, et al. World Wide Web Site-Mouse Imprinting Data and References. Oxfordshire: MRC Hartwell 2013.

41 Tucci V, Isles AR, Kelsey G, Ferguson-Smith AC, Tucci V, Bartolomei MS, et al. Genomic Imprinting and Physiological Processes in Mammals. Cell 2019;176:952–65. 10.1016/j.cell.2019.01.043.

42 Juan AM, Foong YH, Thorvaldsen JL, Lan Y, Leu NA, Rurik JG, et al. Tissue-specific Grb10/Ddc insulator drives allelic architecture for cardiac development. Mol Cell 2022;82:3613–3631.e7. 10.1016/j.molcel.2022.08.021.

43 Wang L, Zhang J, Duan J, Gao X, Zhu W, Lu X, et al. Programming and inheritance of parental DNA methylomes in mammals. Cell 2014;157:979–91. 10.1016/j.cell.2014.04.017.

44 Riemondy KA, Sheridan RM, Gillen A, Yu Y, Bennett CG, Hesselberth JR. valr: Reproducible genome interval analysis in R. F1000Res 2017;6:1025. 10.12688/f1000research.11997.1.

45 Dolinoy DC, Weidman JR, Jirtle RL. Epigenetic gene regulation: linking early developmental environment to adult disease. Reprod Toxicol 2007;23:297–307. 10.1016/j.reprotox.2006.08.012.

46 Thomason ME, Hect JL, Rauh VA, Trentacosta C, Wheelock MD, Eggebrecht AT, et al. Prenatal lead exposure impacts cross-hemispheric and long-range connectivity in the human fetal brain. Neuroimage 2019;191:186–92. 10.1016/j.neuroimage.2019.02.017.

47 Neier K, Montrose L, Chen K, Malloy MA, Jones TR, Svoboda LK, et al. Short- and long-term effects of perinatal phthalate exposures on metabolic pathways in the mouse liver. Environ Epigenet 2020;6:dvaa017. 10.1093/eep/dvaa017.

48 Rygiel CA, Goodrich JM, Solano-González M, Mercado-García A, Hu H, Téllez-Rojo MM, et al. Prenatal Lead (Pb) Exposure and Peripheral Blood DNA Methylation (5mC) and Hydroxymethylation (5hmC) in Mexican Adolescents from the ELEMENT Birth Cohort. Environ Health Perspect 2021;129:67002. 10.1289/EHP8507.

49 Chen C-H, Jiang SS, Chang I-S, Wen H-J, Sun C-W, Wang S-L. Association between fetal exposure to phthalate endocrine disruptor and genome-wide DNA methylation at birth. Environ Res 2018;162:261–70. 10.1016/j.envres.2018.01.009.

50 Sobolewski M, Varma G, Adams B, Anderson DW, Schneider JS, Cory-Slechta DA. Developmental Lead Exposure and Prenatal Stress Result in Sex-Specific Reprograming of Adult Stress Physiology and Epigenetic Profiles in Brain. Toxicol Sci 2018;163:478–89. 10.1093/toxsci/kfy046.

51 Liu S, Wang K, Svoboda LK, Rygiel CA, Neier K, Jones TR, et al. Perinatal DEHP exposure induces sex- and tissue-specific DNA methylation changes in both juvenile and adult mice. Environ Epigenet 2021;7:dvab004. 10.1093/eep/dvab004.

52 Svoboda LK, Ishikawa T, Dolinoy DC. Developmental toxicant exposures and sex-specific effects on epigenetic programming and cardiovascular health across generations. Environ Epigenet 2022;8:dvac017. 10.1093/eep/dvac017.

53 Singh G, Singh V, Sobolewski M, Cory-Slechta DA, Schneider JS. Sex-Dependent Effects of Developmental Lead Exposure on the Brain. Front Genet 2018;9:89. 10.3389/fgene.2018.00089.

54 Smallwood SA, Tomizawa S-I, Krueger F, Ruf N, Carli N, Segonds-Pichon A, et al. Dynamic CpG island methylation landscape in oocytes and preimplantation embryos. Nat Genet 2011;43:811–4. 10.1038/ng.864.

55 Kang E-R, Iqbal K, Tran DA, Rivas GE, Singh P, Pfeifer GP, et al. Effects of endocrine disruptors on imprinted gene expression in the mouse embryo. Epigenetics 2011;6:937–50. 10.4161/epi.6.7.16067.

56 Krishnamoorthy M, Gerwe BA, Scharer CD, Heimburg-Molinaro J, Gregory F, Nash RJ, et al. GABRB3 gene expression increases upon ethanol exposure in human embryonic stem cells. J Recept Signal Transduct Res 2011;31:206–13. 10.3109/10799893.2011.569723.

57 Bastepe M, Fröhlich LF, Hendy GN, Indridason OS, Josse RG, Koshiyama H, et al. Autosomal dominant pseudohypoparathyroidism type Ib is associated with a heterozygous microdeletion that likely disrupts a putative imprinting control element of GNAS. J Clin Invest 2003;112:1255–63. 10.1172/JCI19159.

58 Eggermann T, Begemann M, Kurth I, Elbracht M. Contribution of GRB10 to the prenatal phenotype in Silver-Russell syndrome? Lessons from 7p12 copy number variations. Eur J Med Genet 2019;62:103671. 10.1016/j.ejmg.2019.103671.

59 Wallace C, Smyth DJ, Maisuria-Armer M, Walker NM, Todd JA, Clayton DG. The imprinted DLK1-MEG3 gene region on chromosome 14q32.2 alters susceptibility to type 1 diabetes. Nat Genet 2010;42:68–71. 10.1038/ng.493.

60 Tahara S, Tahara T, Horiguchi N, Okubo M, Terada T, Yoshida D, et al. Lower LINE-1 methylation is associated with promoter hypermethylation and distinct molecular features in gastric cancer. Epigenomics 2019;11:1651–9. 10.2217/epi-2019-0091.

61 Ito Y, Koessler T, Ibrahim AEK, Rai S, Vowler SL, Abu-Amero S, et al. Somatically acquired hypomethylation of IGF2 in breast and colorectal cancer. Hum Mol Genet 2008;17:2633–43. 10.1093/hmg/ddn163.

62 Nye MD, King KE, Darrah TH, Maguire R, Jima DD, Huang Z, et al. Maternal blood lead concentrations, DNA methylation of MEG3 DMR regulating the DLK1/MEG3 imprinted domain and early growth in a multiethnic cohort. Environ Epigenet 2016;2:dvv009. 10.1093/eep/dvv009.

63 Li L, Zhang T, Qin X-S, Ge W, Ma H-G, Sun L-L, et al. Exposure to diethylhexyl phthalate (DEHP) results in a heritable modification of imprint genes DNA methylation in mouse oocytes. Mol Biol Rep 2014;41:1227–35. 10.1007/s11033-013-2967-7.

64 Kochmanski JJ, Marchlewicz EH, Cavalcante RG, Perera BPU, Sartor MA, Dolinoy DC. Longitudinal Effects of Developmental Bisphenol A Exposure on Epigenome-Wide DNA Hydroxymethylation at Imprinted Loci in Mouse Blood. Environmental Health Perspectives n.d.;126:077006. 10.1289/EHP3441.

65 Kalish JM, Jiang C, Bartolomei MS. Epigenetics and imprinting in human disease. Int J Dev Biol 2014;58:291–8. 10.1387/ijdb.140077mb.

66 Prasasya R, Grotheer KV, Siracusa LD, Bartolomei MS. Temple syndrome and Kagami-Ogata syndrome: clinical presentations, genotypes, models and mechanisms. Hum Mol Genet 2020;29:R107–16. 10.1093/hmg/ddaa133.

67 Doshi T, D’souza C, Vanage G. Aberrant DNA methylation at Igf2-H19 imprinting control region in spermatozoa upon neonatal exposure to bisphenol A and its association with post implantation loss. Mol Biol Rep 2013;40:4747–57. 10.1007/s11033-013-2571-x.

68 Schrott R, Greeson KW, King D, Symosko Crow KM, Easley CA, Murphy SK. Cannabis alters DNA methylation at maternally imprinted and autism candidate genes in spermatogenic cells. Syst Biol Reprod Med 2022;68:357–69. 10.1080/19396368.2022.2073292.

69 Soubry A, Hoyo C, Butt CM, Fieuws S, Price TM, Murphy SK, et al. Human exposure to flame-retardants is associated with aberrant DNA methylation at imprinted genes in sperm. Environmental Epigenetics 2017;3:dvx003. 10.1093/eep/dvx003.

70 Weinstein LS, Xie T, Zhang Q-H, Chen M. Studies of the regulation and function of the Gsα gene Gnas using gene targeting technology. Pharmacol Ther 2007;115:271–91. 10.1016/j.pharmthera.2007.03.013.

71 Wang L, Chang S, Wang Z, Wang S, Huo J, Ding G, et al. Altered GNAS imprinting due to folic acid deficiency contributes to poor embryo development and may lead to neural tube defects. Oncotarget 2017;8:110797–810. 10.18632/oncotarget.22731.

72 Hanna P, Francou B, Delemer B, Jüppner H, Linglart A. A Novel Familial PHP1B Variant With Incomplete Loss of Methylation at GNAS-A/B and Enhanced Methylation at GNAS-AS2. J Clin Endocrinol Metab 2021;106:2779–87. 10.1210/clinem/dgab136.

73 Turan S, Bastepe M. The GNAS complex locus and human diseases associated with loss-of-function mutations or epimutations within this imprinted gene. Horm Res Paediatr 2013;80:10.1159/000355384. 10.1159/000355384.

74 Wroe SF, Kelsey G, Skinner JA, Bodle D, Ball ST, Beechey CV, et al. An imprinted transcript, antisense to Nesp, adds complexity to the cluster of imprinted genes at the mouse Gnas locus. Proc Natl Acad Sci U S A 2000;97:3342–6. 10.1073/pnas.97.7.3342.

75 Hayward BE, Kamiya M, Strain L, Moran V, Campbell R, Hayashizaki Y, et al. The human GNAS1 gene is imprinted and encodes distinct paternally and biallelically expressed G proteins. Proc Natl Acad Sci U S A 1998;95:10038–43. 10.1073/pnas.95.17.10038.

76 Desbuquois B, Carré N, Burnol A-F. Regulation of insulin and type 1 insulin-like growth factor signaling and action by the Grb10/14 and SH2B1/B2 adaptor proteins. FEBS J 2013;280:794–816. 10.1111/febs.12080.

77 Plasschaert RN, Bartolomei MS. Tissue-specific regulation and function of Grb10 during growth and neuronal commitment. Proc Natl Acad Sci U S A 2015;112:6841–7. 10.1073/pnas.1411254111.

78 Hikichi T, Kohda T, Kaneko-Ishino T, Ishino F. Imprinting regulation of the murine Meg1/Grb10 and human GRB10 genes; roles of brain-specific promoters and mouse-specific CTCF-binding sites. Nucleic Acids Res 2003;31:1398–406. 10.1093/nar/gkg232.

79 Luo L, Jiang W, Liu H, Bu J, Tang P, Du C, et al. De-silencing Grb10 contributes to acute ER stress-induced steatosis in mouse liver. J Mol Endocrinol 2018;60:285–97. 10.1530/JME-18-0018.

80 Blagitko N, Mergenthaler S, Schulz U, Wollmann HA, Craigen W, Eggermann T, et al. Human GRB10 is imprinted and expressed from the paternal and maternal allele in a highly tissue- and isoform-specific fashion. Human Molecular Genetics 2000;9:1587–95. 10.1093/hmg/9.11.1587.

81 Bakulski KM, Feinberg JI, Andrews SV, Yang J, Brown S, L. McKenney S, et al. DNA methylation of cord blood cell types: Applications for mixed cell birth studies. Epigenetics 2016;11:354–62. 10.1080/15592294.2016.1161875.

82 Armand EJ, Li J, Xie F, Luo C, Mukamel EA. Single-Cell Sequencing of Brain Cell Transcriptomes and Epigenomes. Neuron 2021;109:11–26. 10.1016/j.neuron.2020.12.010.

83 Campbell KA, Colacino JA, Park SK, Bakulski KM. Cell types in environmental epigenetic studies: Biological and epidemiological frameworks. Curr Environ Health Rep 2020;7:185–97. 10.1007/s40572-020-00287-0.

