## Supplementary Figures for "Effects of Developmental Lead and Phthalate Exposures on DNA Methylation in Adult Mouse Blood, Brain, and Liver Identifies Tissue- and Sex-Specific Changes with Implications for Genomic Imprinting"

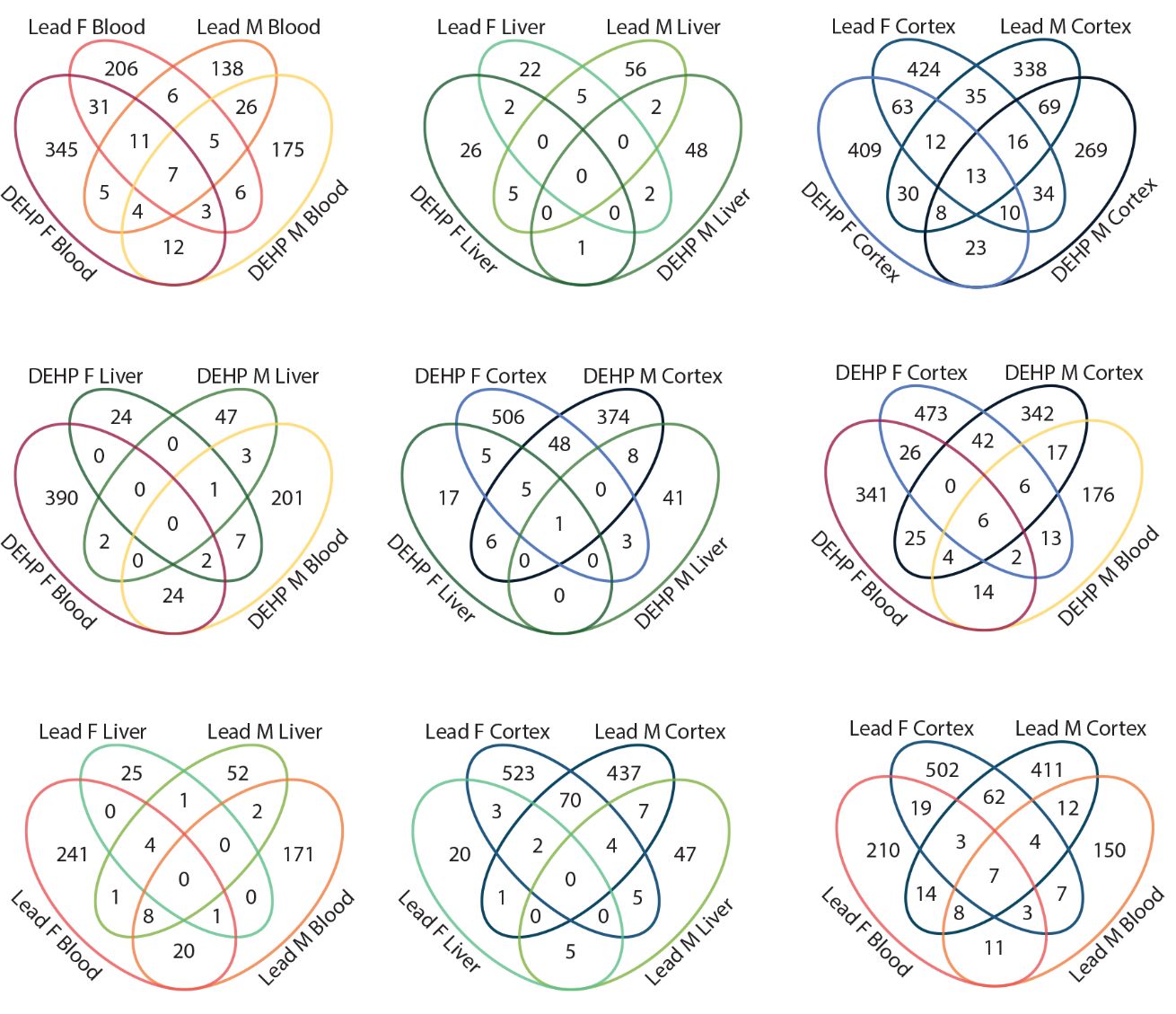

Supplementary Figure 1. DMR containing genes across different tissues, sex, and exposure groups.

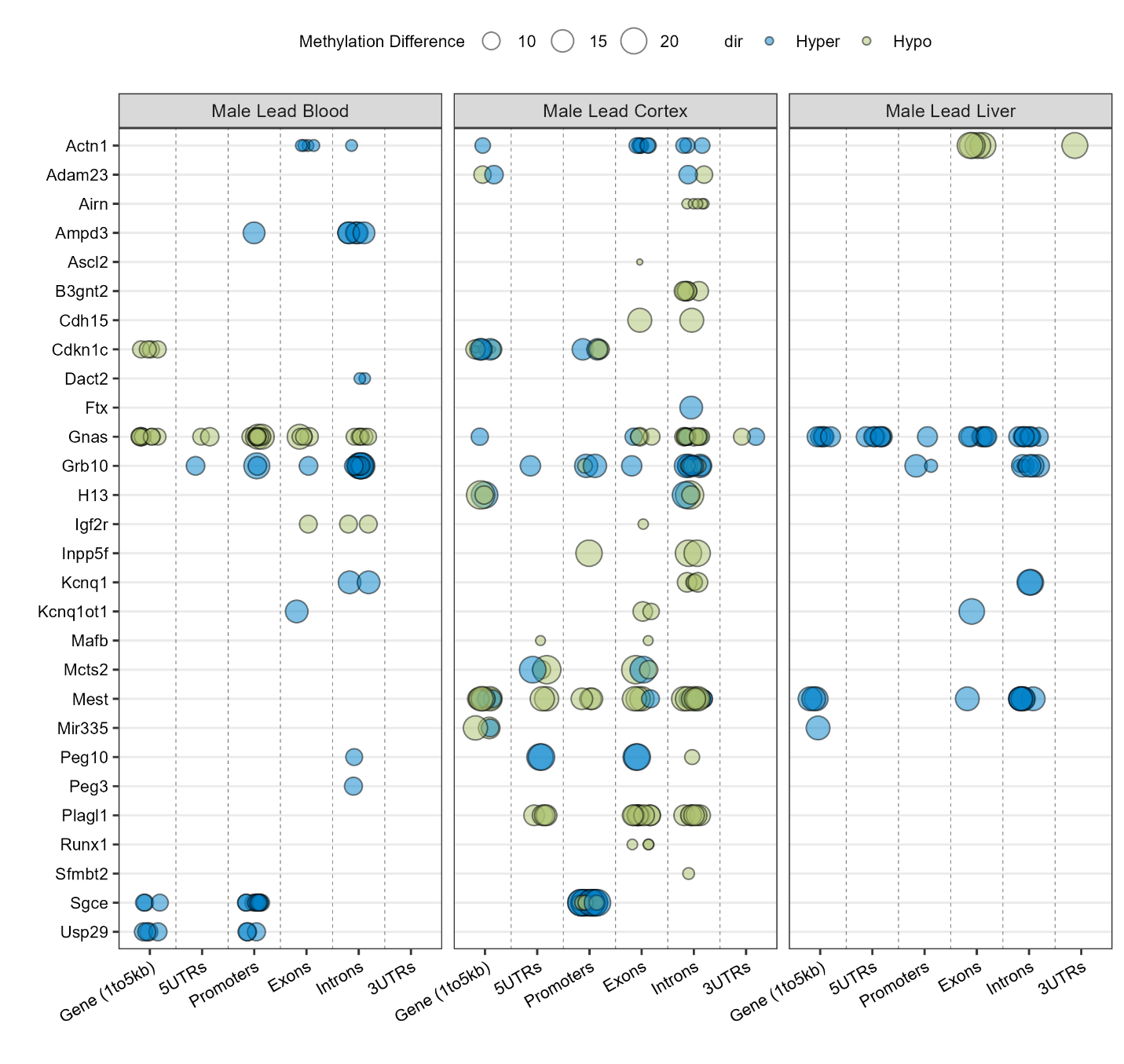

Supplementary Figure 2. DMR containing imprinted genes in male tissues with lead exposure.

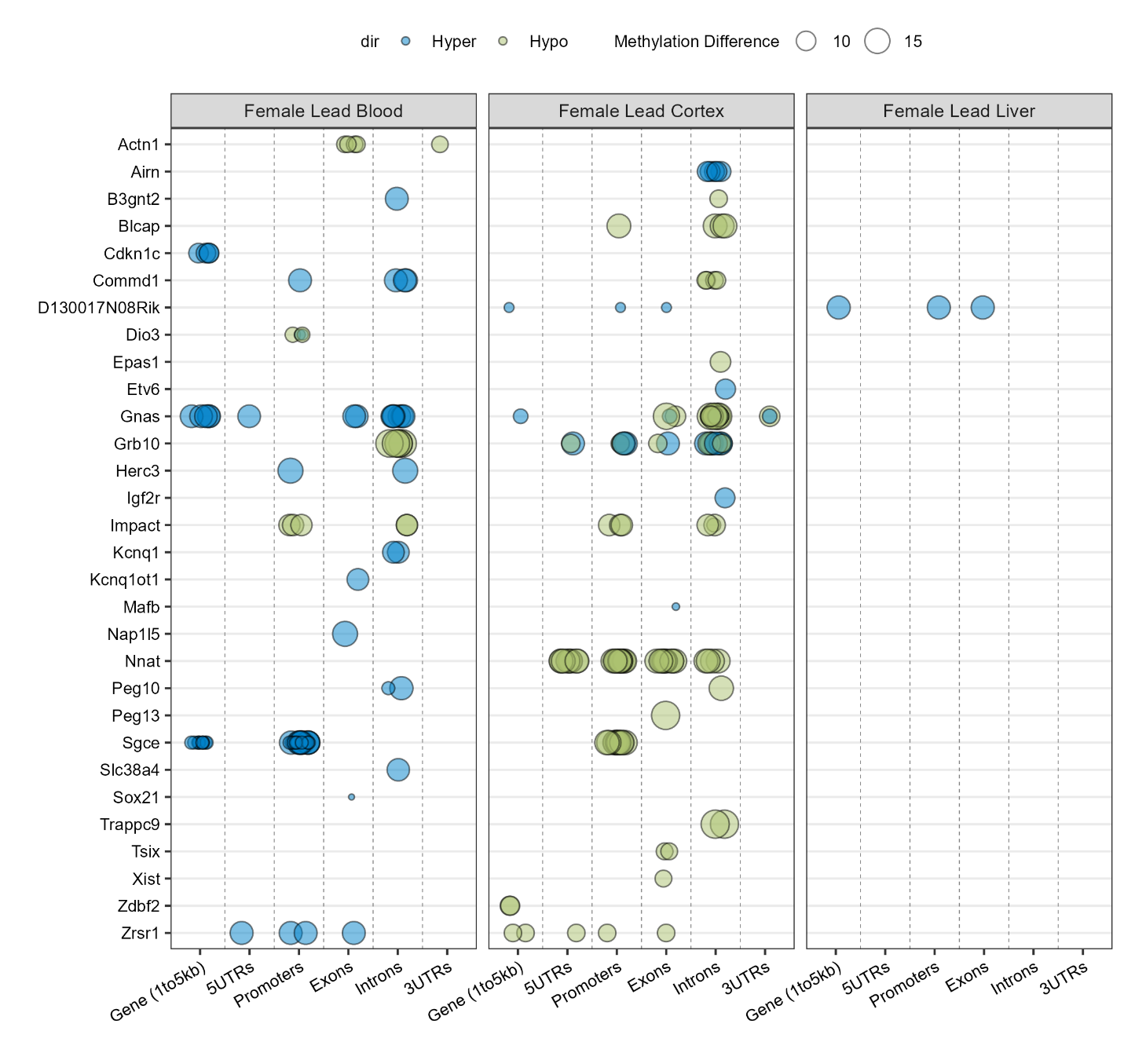

Supplementary Figure 3. DMR containing imprinted genes in female tissues with lead exposure.

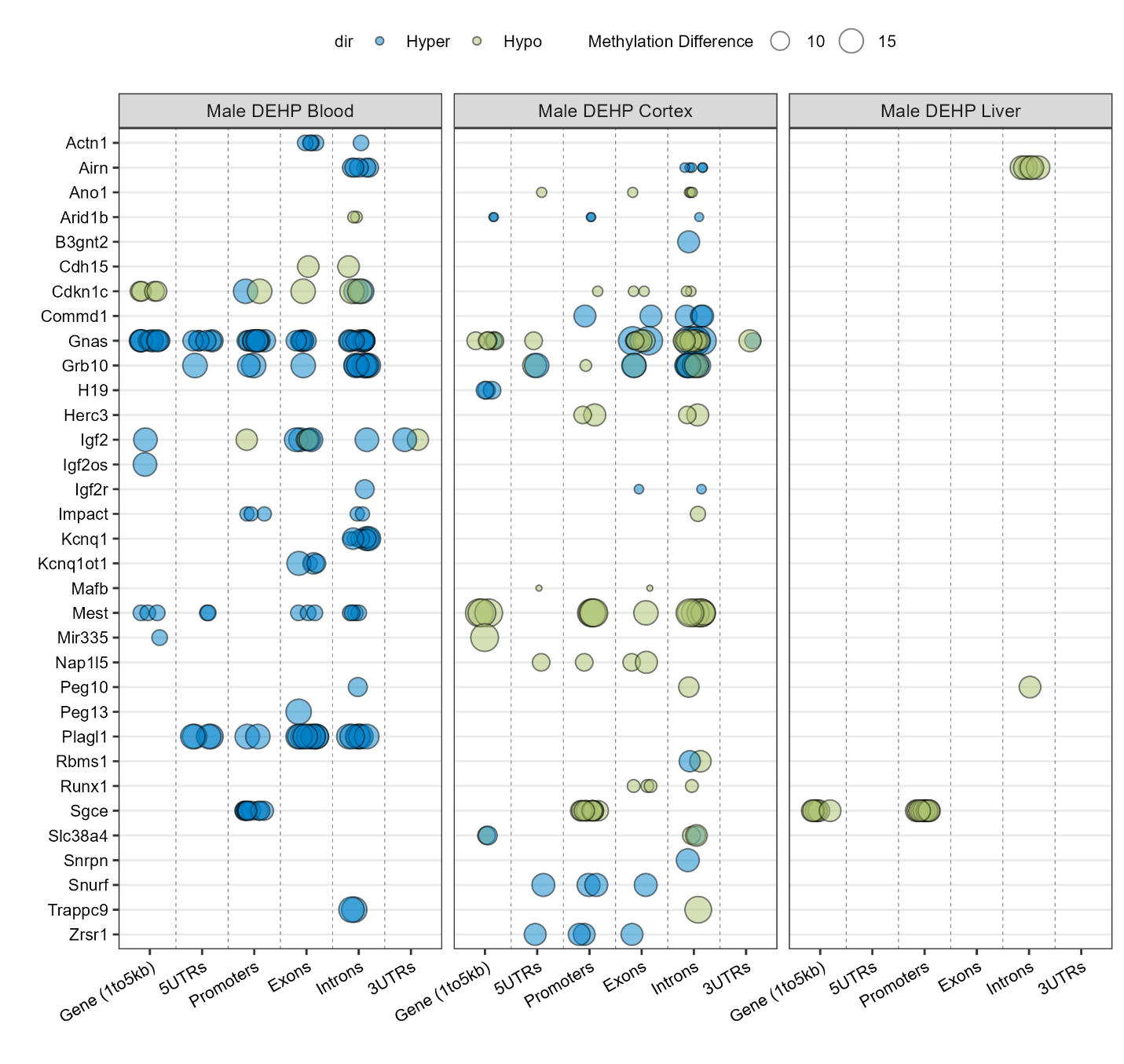

Supplementary Figure 4. DMR containing imprinted genes in male tissues with DEHP exposure.

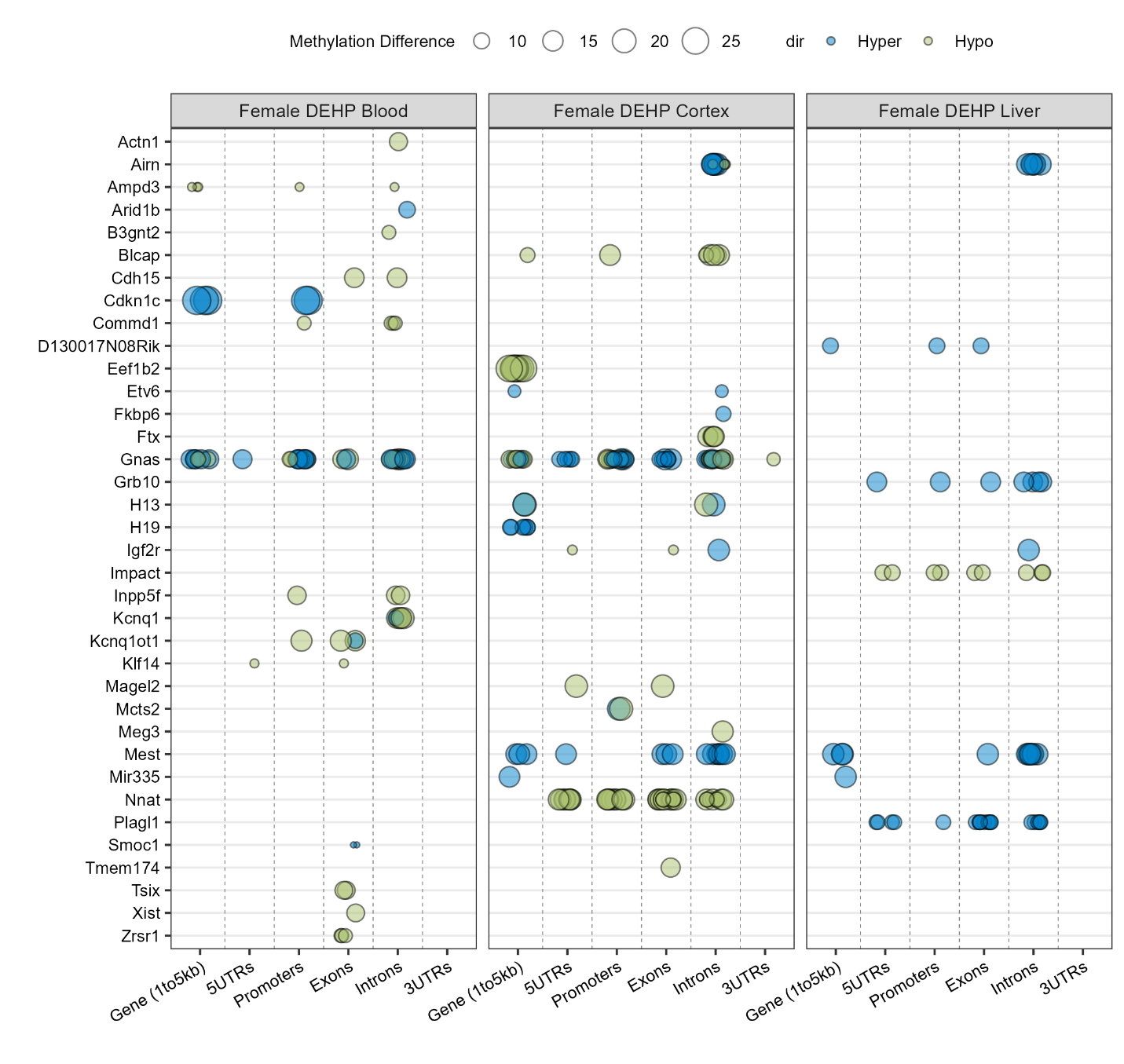

Supplementary Figure 5. DMR containing imprinted genes in female tissues with DEHP exposure.

Supplementary Table 1. Number of DMRs detected by MethylSig and Metilene.

| Exposure | Sex | Tissue | Method | | Overlap DMRs | After merge DMRs |
| --- | --- | --- | --- | --- | --- | --- |
|  |  |  | MethylSig | Metilene |  |  |
| DEHP | Male | Blood | 146 | 173 | 7 | 312 |
|  |  | Liver | 81 | 11 | 1 | 90 |
|  |  | Cortex | 195 | 417 | 25 | 587 |
|  | Female | Blood | 131 | 355 | 9 | 477 |
|  |  | Liver | 16 | 25 | 1 | 40 |
|  |  | Cortex | 189 | 495 | 24 | 661 |
| Lead | Male | Blood | 93 | 153 | 3 | 243 |
|  |  | Liver | 77 | 24 | 1 | 100 |
|  |  | Cortex | 220 | 500 | 32 | 688 |
|  | Female | Blood | 58 | 241 | 7 | 292 |
|  |  | Liver | 23 | 15 | 2 | 36 |
|  |  | Cortex | 209 | 567 | 31 | 746 |

Supplementary Table 2. Percentage of genomic annotation for detected DMRs.

Supplementary Table 3. Number of DMR-containing genes associated with Go terms.

Supplementary Table 4. DMR associated mouse imprinted genes.

Supplementary Table 5. Binomial test of DMRs located in imprinted control regions.

Supplementary Table 6. Lead exposure DMRs associated with mouse imprinted gene control regions.

Supplementary Table 7. DEHP exposure DMRs associated with mouse imprinted gene control regions.
